# CRISPR base editing of *cis*-regulatory elements enables target gene perturbations

**DOI:** 10.1101/2022.05.26.493649

**Authors:** Colin K.W. Lim, Tristan X. McCallister, Christian Saporito-Magriña, Garrett D. McPheron, Ramya Krishnan, M. Alejandra Zeballos C, Jackson E. Powell, Lindsay V. Clark, Pablo Perez-Pinera, Thomas Gaj

## Abstract

CRISPR technology has demonstrated broad utility for controlling target gene expression; however, there remains a need for strategies capable of modulating expression via the precise editing of non-coding regulatory elements. Here we demonstrate that CRISPR base editors, a class of gene-modifying proteins capable of creating single-base substitutions in DNA, can be used to perturb gene expression via their targeted mutagenesis of *cis*-acting sequences. Using the promoter region of the human huntingtin (HTT) gene as an initial target, we show that editing of the binding site for the transcription factor NF-κB led to a marked reduction in HTT gene expression in base-edited cell populations. We found that these gene perturbations were persistent and specific, as a transcriptome-wide RNA analysis revealed minimal off-target effects resulting from the action of the base editor protein. We further demonstrate that this base-editing platform could influence gene expression *in vivo* as its delivery to a mouse model of Huntington’s disease led to a potent decrease in HTT mRNA in striatal neurons. Finally, to illustrate the applicability of this concept, we target the amyloid precursor protein, showing that multiplex editing of its promoter region significantly perturbed its expression. These findings demonstrate the potential for base editors to regulate target gene expression.

## INTRODUCTION

Gene expression, the process by which information encoded within DNA is converted to a functional product, is a highly controlled process involving the coordinated action of transcription factors (TFs) that bind to *cis*-regulatory elements to orchestrate transcription.^1-3^ Given its importance for maintaining cell homeostasis during development and adulthood, as well as the role that its dysregulation can play in disease, considerable effort has been devoted towards the development of strategies capable of manipulating target gene expression.^4,5^ Among these are approaches based on Cas9,^6^ an RNA-guided DNA endonuclease that, since its discovery as a programmable nuclease involved in adaptive immunity in bacteria,^6-9^ has been repurposed for targeted gene editing in eukaryotic cells.^6,10-12^ Cas9 can be directed to a specific DNA site by a single guide RNA (sgRNA) that encodes a programmable spacer sequence that mediates DNA binding via RNA-DNA base complementarity.^6^ Once bound to its target, Cas9 cleaves it,^13^ activating DNA repair pathways such as non-homologous end joining (NHEJ) which can introduce random base insertions and deletions (indels) at the target site.^10,11,14^ This outcome, in particular, can be exploited to disrupt *cis*-acting sequences, which in turn can influence the expression of a target gene.^15-17^ Catalytically inactive forms of Cas9, however, can also be used to regulate expression,^18-20^ as dead Cas9 (dCas9) can be deployed to physically block transcriptional initiation and/or elongation of a target transcript,^18,20^ or epigenetically silence or activate the expression of an endogenous locus when tethered to a transcriptional repressor,^19,21^ or activator domain.^22-26^

However, while effective, these approaches possess several limitations that could restrict their utility for certain applications. For instance, to sustain their effectiveness, CRISPR repressors and activators must be persistently expressed in cells in order to continuously engage with their target sequence(s), a technical limitation that could increase their likelihood for affecting the expression of non-target genes. Additionally, Cas9-introduced indels, which can be directed to *cis*-acting elements to perturb expression, are heterogenous, can vary in size,^27-29^ and pose risk for destroying TF binding sites,^15-17^ which may be undesirable for applications that require the fine-tuning of target gene expression. Finally, while alternative pathways such as homology-directed repair (HDR) can be used to precisely edit bases in *cis*-acting elements,^10,11^ HDR is not available at high rates in many cell types,^10,30-32^ which can limit its implementation. As a result, there remains a need for strategies capable of modulating target gene expression via the precise editing of *cis*-acting sequences using pathways available within a broad range of cell types.

A class of genome-modifying technologies with the potential to meet these needs are CRISPR base editors.^33^ Generally consisting of fusions of a targetable Cas9 nickase (nCas9) variant with a nucleobase deaminase enzyme,^34,35^ base editors have the ability to induce targeted single-base substitutions in DNA, but without the need for a DSB, thereby overcoming a disadvantage inherent to NHEJ-based approaches for DNA editing. In particular, unlike Cas9 nucleases, base editors have a defined catalytic window that can enable the selective editing of a target base within a *cis*-regulatory sequence. Base editors also yield predictable editing outcomes, a feature that could be exploited to enable reproducible changes in target gene expression. Given these attributes, we hypothesized that base editors could be an attractive platform for modulating target gene expression via their precise editing of non-coding regulatory elements.

Here, we demonstrate that mutagenizing *cis*-acting sequences by base editing provides a means for perturbing target gene expression. Following the identification of actionable elements in the promoter region of the human huntingtin (HTT) gene, which we initially targeted to validate this concept, we show that base editing the binding site for the transcription factor (TF) NF-κB led to a marked decrease in HTT expression in base-edited cell populations. We found that editing of the HTT promoter was stable, resulting in a persistent decrease in HTT mRNA and protein over time, and specific, as RNA-seq revealed minimal differentially expressed genes from base editing. We further show that this particular base-editing platform could lower HTT *in vivo*, as its intrastriatal delivery to a mouse model of Huntington’s disease (HD) led to a reduction in HTT gene expression within neurons. Finally, we illustrate the applicability of this approach by targeting the amyloid precursor protein, demonstrating that multiplexed editing of its promoter region could significantly perturb its expression. Thus, our findings demonstrate the ability for base editors to regulate target gene expression.

## RESULTS

### Identification of actionable elements in the HTT promoter

We sought to determine the ability for base editors to influence the expression of target gene(s) via their targeted mutagenesis of *cis*-regulatory sequences. As an initial proof of concept, we targeted the promoter region for the HTT gene, which encodes a protein that, when mutated to carry >35 copies of a CAG trinucleotide within exon 1 of its coding sequence, causes HD, a debilitating and ultimately fatal neurodegenerative disorder characterized by the loss of neurons in the striatum.^36^

There exists in the human HTT promoter predicted binding sites for the TFs NF-κB (-139 bps from the translation start codon), AP2 (-242), SP1 (-280), AP4 (-326) and AML1 (-392) (**Figure 1A**).^37,38^ Given the number of candidate base editors that could be created to span these regions, as well as the prospect of other important regulatory elements within it, we sought to streamline our design efforts by creating a map of functional elements for a portion of the HTT promoter. To this end, we conducted a CRISPR interference (CRISPRi) screen using 30 sgRNAs designed to tile the human HTT promoter from -700 to -30 bps from the translation start codon (**Figure 1A**), reasoning that such a screen would enable the identification and/or ranking of sequences most suitable for targeting by base editing. To facilitate the identification of sgRNAs that could perturb expression, we created a reporter plasmid expressing a Renilla luciferase transgene from a ∼1 kb fragment of the human HTT promoter (pHTT-RLuc), with the expectation that dCas9 binding to functional elements within it would alter Renilla luciferase expression.

**Figure 1.**
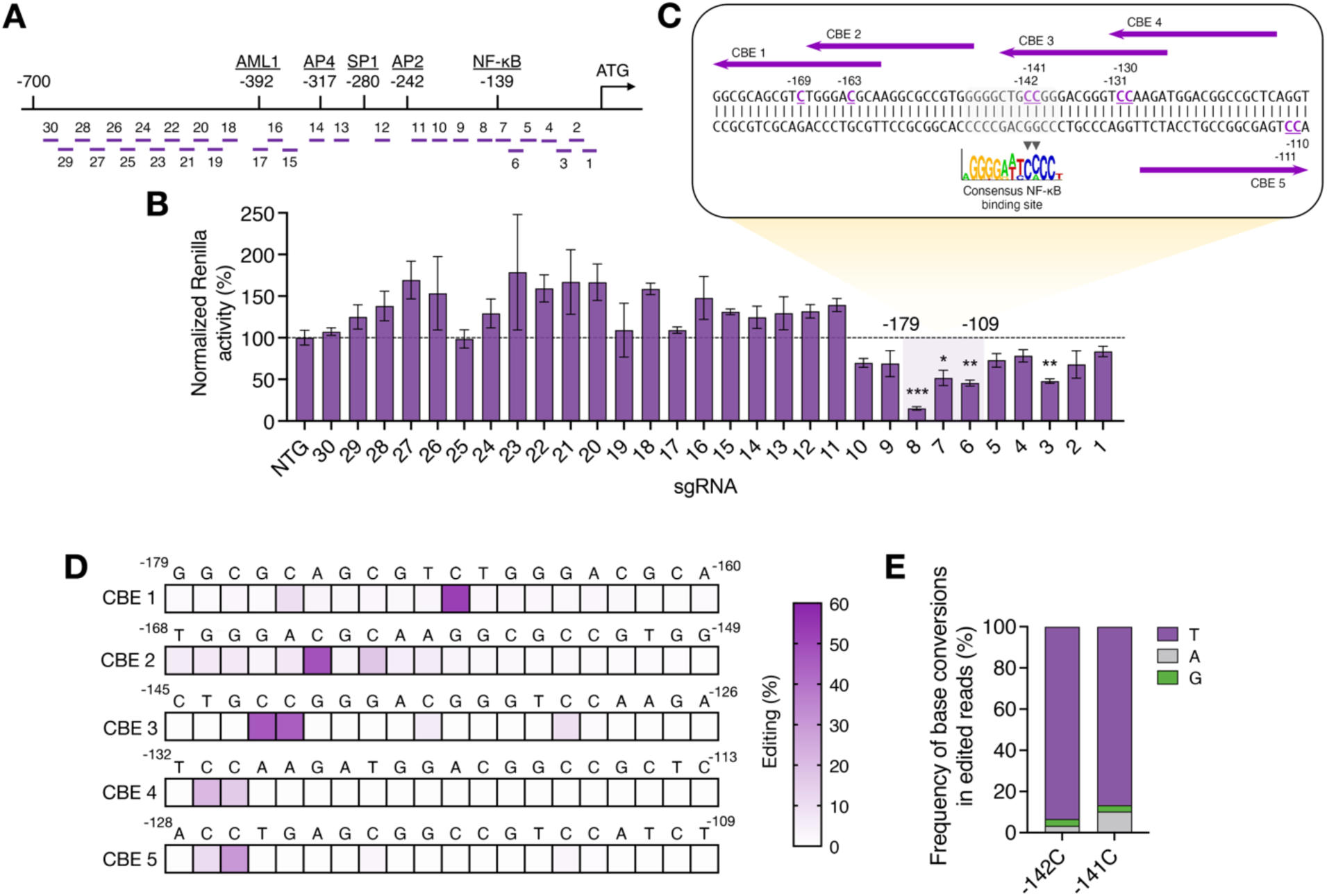
CRISPR interference can identify actionable elements in the human HTT promoter. **(A)** Cartoon showing the positions of the binding sites for the transcription factors (TFs) NF-κB, AP2, SP1, AP4 and AML1 in the promoter region of the human HTT gene. The approximate locations of the sgRNA binding sites for the CRISPR interference (CRISPRi) screen are indicated by purple bars. **(B)** Normalized Renilla luciferase expression in HEK293T cells after transfection with the pHTT-RLuc reporter and expression vectors encoding dCas9 and each sgRNA. Renilla expression in each transfection was normalized to firefly luciferase, and all relative values were normalized to those from cells trasfected with pHTT-RLuc and dCas9 with a non-targeted sgRNA. **(C)** Sequence of the human HTT promoter and the NF-κB binding site, with the CBE binding sites indicated by purple arrows. Underlined bases denote the target cytosines for each CBE, with the numbering specifying their position to the translation start site. The NF-κB consensus motif is shown in the logo cartoon ^45^, with black arrowheads indicating the target cytosines for CBE-3. **(D)** Heat map showing the mean editing frequency for each base by the candidate CBEs in HEK293T cells, as determined by deep sequencing (n = 3). Numbering indicates the relative position of the base to the translation start codon. **(E)** Base conversion frequencies within edited reads at positions -142C and -141C in the HTT promoter in HEK293T cells transfected with CBE-3, as determined by deep sequencing (n = 3). Bars indicate means and error bars indicate S.E.M. *P < 0.05, **P < 0.01, ***P < 0.001; one-tailed unpaired t-test.

To implement this screen, we transfected human embryonic kidney (HEK) 293T cells with expression vectors encoding dCas9 and each of the 30 sgRNAs tiling the HTT promoter in combination with pHTT-RLuc, a plasmid that also contained within it the coding sequence for firefly luciferase (which we used to normalize for transfection efficiency). From this screen, we found that sgRNAs targeting -179 to -110 bps from the translation start codon, a region that contains near its center the predicted binding site for NF-κB, effectively repressed Renilla expression (**Figure 1B**). The most active of these sgRNAs, in particular, targeted the NF-κB binding site and reduced Renilla activity by ∼85% (P < 0.001; **Figure 1B**), which is consistent with prior studies demonstrating that NF-κB can modulate HTT expression.^38,39^ Additionally, we found that an sgRNA targeting a sequence ∼46 bps from the translation start codon also effectively repressed Renilla expression (P < 0.01; **Figure 1B**); however, unlike the sgRNAs that targeted the regions flanking the NF-κB binding site, the sgRNAs targeting the positions adjacent to this site had no significant effect on Renilla (P > 0.05; **Figure 1B**). Interestingly, sgRNAs targeting ±10 bps of the predicted binding sites for the TFs Sp1, AP2, AP4, and AML1, as well as the more distal regions of the cloned promoter fragment (-700 to -400 bps) were all found to either have no effect Renilla expression or increased it (**Figure 1B**), indicating their poor suitability as candidates for targeting by base editing.

Overall, our results demonstrate that the NF-κB binding site and the sequences surrounding it are important for modulating expression from the HTT promoter.

### Base editing of the NF-κB binding site in the HTT promoter can decrease gene expression

Next, we sought to capitalize on the information from the CRISPRi screen by determining whether base editing of the NF-κB binding site and/or the sequences adjacent to it could regulate HTT gene expression. Because of the GC-rich nature of this 70-bp window, we targeted it exclusively with cytosine base editors (CBEs),^34,40,41^ a class of base-editing protein that relies on a cytidine deaminase domain (in this case the catalytic domain from the enzyme APOBEC1) to induce the deamination of cytosines, which then facilitates their conversion to thymines. More specifically, we designed five CBEs to tile this 70-bp region with the expectation that C > T editing within this window could disrupt the binding of a *cis*-acting regulator that positively controls HTT expression (**Figure 1C**). Notably, several of the CBEs that we designed were expected to target multiple cytosines, including CBE-3, a variant whose editing window overlaps with the predicted binding site for NF-κB and includes two conserved cytosines (-142C and -141C) (**Figure 1C)**.

We first measured the ability of the designed CBEs to edit the HTT promoter in HEK293T cells using deep sequencing. Of the five CBEs tested, we found that three variants, CBE-1 and CBE-2 (which target sequences 5’ of the NF-κB binding site) and CBE-3 (which targets the NF-κB binding site) edited their target cytosines with efficiencies >25% (P < 0.001; **Figure 1D**). CBE-3, in particular, was found to edit its two target bases (-142C and -141C) with efficiencies near 40% (P < 0.001; **Figure 1D)** with deep sequencing revealing that ∼90% and ∼85% of these base conversions encoded the target thymine (**Figure 1E**).

Given its higher editing efficiency compared to the other variants and the predicted importance of positions -142C and -141C for NF-κB binding (**Figure 1C**), we next tested the ability of CBE-3 to modulate the expression of HTT. To restrict our analysis to base-edited cells rather than a bulk population consisting of edited and non-edited cells that could dampen the measured effect,^42,43^ we co-delivered CBE-3 with a transient reporter that encodes a blue fluorescent protein (BFP) variant that can be converted to a green fluorescent protein (GFP) following the induction of a C > T edit driven by a co-transfected BFP-targeting sgRNA,^42^ enabling the enrichment of base-edited cells by fluorescence-activated cell sorting (FACS).

As expected, transient enrichment increased the proportion of cells with base edits in the NF-κB binding site, as ∼65% of the analyzed alleles from the enriched pool of cells harbored edits at -142C and -141C (**Figure S1**). Using qPCR, we measured the expression of the HTT gene in the enriched cell populations. Compared to similarly enriched cells that were transfected with a non-targeting CBE, we observed a ∼60% decrease in HTT mRNA in cells transfected with CBE-3 (P < 0.001) (**Figure 2A**).

**Figure 2.**
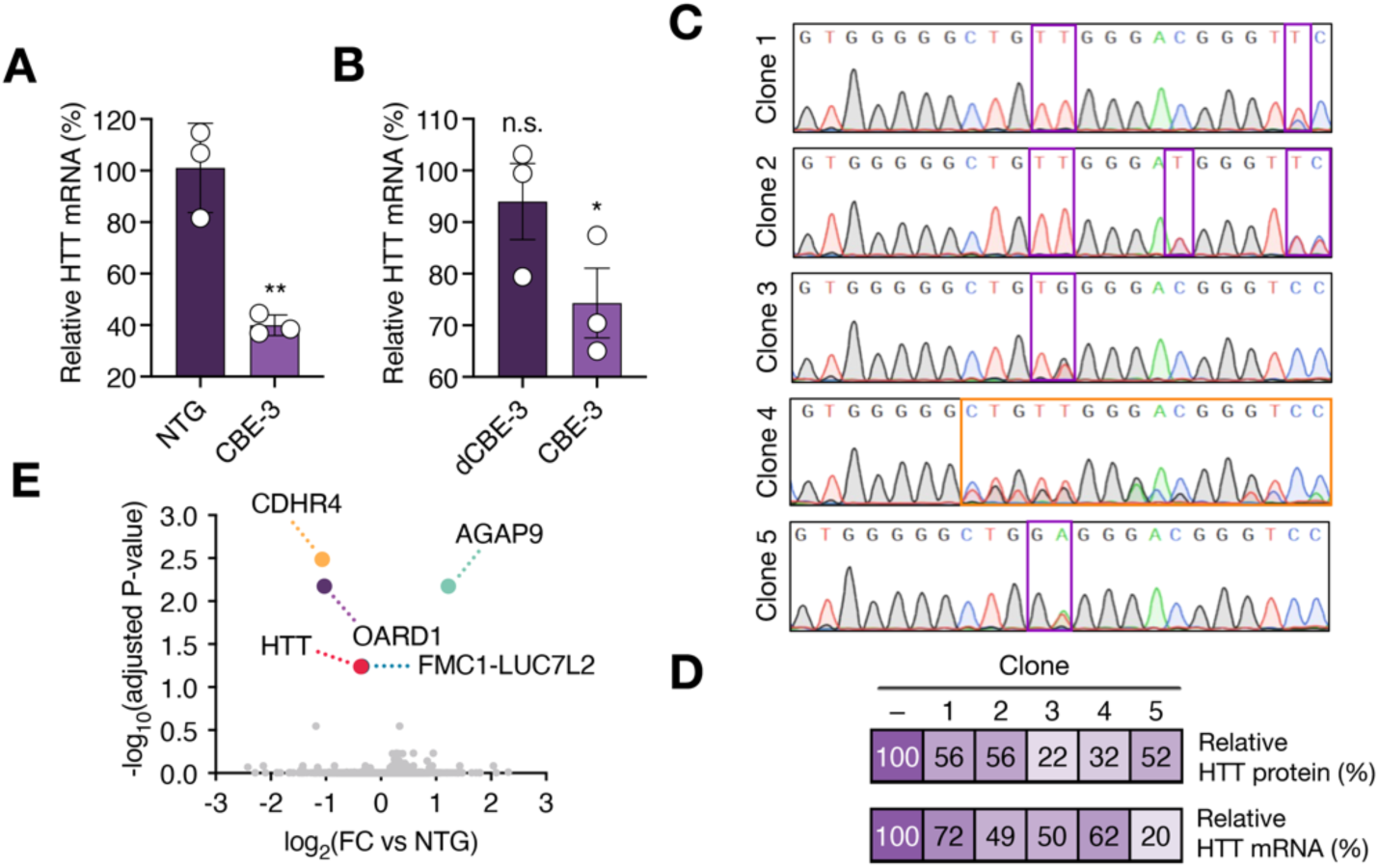
Base editing of the NF-κB binding site can lower HTT expression. **(A)** Relative HTT mRNA in HEK293T cells transfected with a targeted or non-targeted CBE and enriched by FACS using a transient reporter (n = 3). **(B)** Relative HTT mRNA in HEK293T cells seven days after transfection with the CBE-3 or a deactivated CBE-3 (dCBE-3). (**A** and **B**) All data were normalized to HTT mRNA in cells transfected with a non-targeted CBE (n = 3). **(C)** Sanger sequencing of the NF-κB binding site in expanded HEK293T clones originally transfected with CBE-3. Purple boxes indicate the bases edited by CBE-3, while the orange box indicates an indel mutation. **(D)** Heat map showing (top) relative HTT protein and (bottom) relative HTT mRNA in the expanded HEK293T clones. Values are relative to the measured HTT protein and mRNA from unedited HEK293T clones. **(E)** Volcano plot of RNA-seq data comparing HEK293T cells transfected with CBE-3 to cells transfected with a non-targeted CBE (n = 3). The red circle indicates the HTT gene, while the colored circles indicate non-target differentially expressed genes [>1.25-fold change (FC), FDR-adjusted P < 0.1]. Bars represent means and error bars indicate S.E.M. *P < 0.05, **P <; one-tailed unpaired t-test.

However, because dCas9 binding to a promoter region can interfere with transcriptional initiation and decrease target gene expression,^18^ we next sought to determine if the base-editing outcomes themselves – and not interference from the binding of the CBE to its target – were driving the change in HTT expression. To this end, we created a deactivated CBE (dCBE) variant containing inactivating mutations (H61A and E63A) in its APOBEC1 catalytic domain,^44^ but which still possessed the ability to bind DNA (**Figure S2**). Using qPCR, we measured the ability of the dCBE targeted to the NF-κB binding site (dCBE-3) to lower HTT expression in HEK293T cells at seven days post-transfection, a time point that we reasoned would provide sufficient time for the edits in the HTT promoter to impact transcription of the HTT gene. From this analysis, we found that only cells transfected with the catalytically active form of the CBE had reduced HTT expression (P < 0.05) (**Figure 2B**), as we measured no significant difference in HTT mRNA in cells transfected with dCBE-3 versus a non-targeted CBE (P > 0.05) (**Figure 2B**).

To further establish that base-editing of the NF-κB binding site in the HTT promoter reduced HTT expression, we analyzed single cell-derived HEK293T clones that originated from a bulk pool of cells transfected with CBE-3 and contained the target -142C > T and -141C > T edits (**Figure 2C**) (further, though found to occur at low frequencies, we expanded clones with non-target C > A and C > G edits, as well as indels, to determine the effect that non-precise editing outcomes at -142C and -141C could have on HTT expression). We measured by western blot a nearly two-fold decrease in HTT protein for each edited clone compared to a control cell line (**Figure 2D**), with no CBE protein detected in the lysate of any clone (**Figure S3**), indicating that the decrease in HTT was attributable to the base edits. These results were further corroborated by qPCR, which, depending on the clone, revealed a ∼30-50% reduction in HTT mRNA compared to the control cell lines (**Figure 2D**). Interestingly, though unsurprisingly, our analyses revealed that clones harboring non-target edits had a greater reduction in HTT mRNA and protein compared to clones with the target C > T edits, a finding that is consistent with the NF-κB consensus motif, which predicts that C > G substitutions at -142C and -141C should be particularly disruptive (**Figure 1C**).^45^

Using RNA-seq, we next determined if the action of the CBE led to differential gene expression changes in HEK293T cells. In addition to the HTT gene, whose downregulation was anticipated, we measured that only four non-target genes were differentially expressed in cells transfected with CBE-3 compared to cells transfected with a non-targeted CBE (>1.25-fold change; false discovery rate (FDR)–adjusted P < 0.1) (**Figure 2E and Supplementary Table S2**).

Of the four differentially expressed genes (DEGs), we found that none of them contained pseudo-sgRNA binding sites (which we defined as having >75% of contiguous identity to the CBE-3 target) within 5 kb upstream or downstream of their respective transcription start site. An overrepresentation analysis of gene ontology terms further revealed no significant enrichment for any themes among HTT and the DEGs, indicating an unknown mechanism of dysregulation for the four non-target genes.

In addition to determining its transcriptome-wide effects, we evaluated if the CBE edited off-target sites in the human genome. Using deep sequencing, we measured no editing at any of the analyzed computationally predicted off-target sites for CBE-3 in HEK293T cells (P > 0.1; **Figure S4**).

Collectively, these results demonstrate that base editing of the NF-κB binding site in the promoter region of the human HTT gene can perturb its expression. Additionally, we show that the action of this base editor led to minimal changes in the transcriptome of cells.

### *In vivo* targeting of the HTT promoter by base editing can modulate HTT expression

We next sought to determine whether the base editing system designed to target the HTT promoter could reduce HTT expression *in vivo*. More specifically, we sought to evaluate the targeting capabilities of CBE-3 in the R6/2 mouse model of HD, which carries a ∼1 kb fragment of the human HTT promoter that drives expression of the toxic N-terminal fragment of a mutant HTT (mHTT) protein with ∼120 polyQ repeats.^46^ R6/2 mice develop a progressive phenotype that involves the accumulation of mHTT inclusions in striatal neurons and is characterized in part by a shortened lifespan.^46-48^

To deliver the CBE *in vivo*, we used adeno-associated virus (AAV). Specifically, we used AAV1, which is capable of transducing neurons following its direct injection to the striatal parenchyma.^49-54^ However, because AAV possesses a limited carrying capacity that restricts its ability to deliver a full-length CBE by a single vector, we employed a split-intein-containing CBE scaffold that we,^55,56^ and others^57-59^ have demonstrated is compatible with dual vector delivery and can be used to reconstitute a functional, full-length CBE protein to enable *in vivo* DNA editing (**Figure 3A**). Thus, we intra-striatally injected four-week-old R6/2 mice with ∼3 × 10^10^ viral particles each of two AAV1 vectors encoding either the N- or C-terminal split-intein CBE domains with sgRNAs targeting the human HTT promoter (AAV1-CBE-hHTT) or the mouse Rosa26 locus (AAV1-CBE-mRosa26) (**Figure 3B**). Additionally, we co-injected R6/2 mice with ∼3 × 10^10^ viral particles of a third AAV1 vector encoding an EGFP variant fused to a KASH (Klarsicht/ANC-1/Syne-1 homology) domain (AAV1-EGFP-KASH) (**Figure 3B**). EGFP-KASH localizes to the outer nuclear membrane, enabling the isolation of transduced neuronal nuclei by FACS for a higher resolution analysis of base editor-mediated outcomes.^57,60,61^

**Figure 3.**
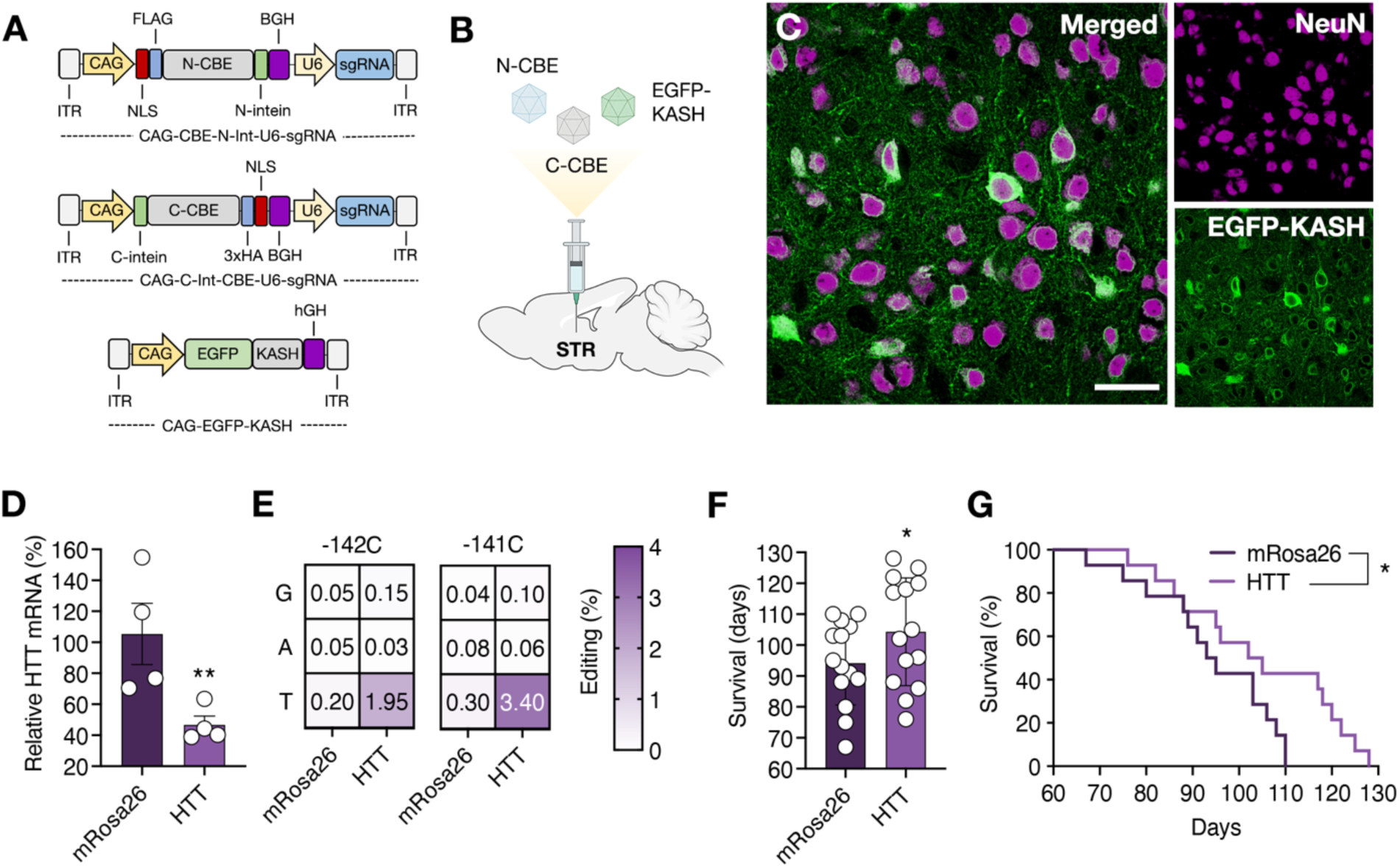
Targeting the HTT promoter *in vivo* can lower HTT expression in a mouse model of HD. **(A)** Schematic of the AAV vectors used in this study. Abbreviations are as follows: ITR, inverted terminal repeat; CAG, cytomegalovirus early enhancer/chicken β-actin promoter; NLS, nuclear localization signal; FLAG, FLAG epitope tag; 3x HA, three repeats of the human influenza hemagglutinin (HA) epitope tag. **(B)** Overview of the intra-striatal injections conducted on R6/2 mice. **(C)** Representative immunofluorescence staining of the striatum four weeks after mice were injected with 3 × 10^10^ particles each of dual AAV1 particles encoding the N- or C-terminal split-intein CBE domains with sgRNAs targeting the human HTT promoter (AAV1-CBE-hHTT) or the mouse Rosa26 locus (AAV1-CBE-mRosa26) and 3 × 10^10^ particles of AAV1-EGFP-KASH. Scale bar, 30 μm. **(D)** Relative HTT mRNA (n = 4) and (**E**) heat map showing the mean editing frequencies at positions -142C and -141C in the human HTT promoter (n = 3) in FACS-enriched EGFP^+^ nuclei from treated (AAV1-CBE-HTT) or untreated (AAV1-CBE-mRosa26) R6/2 mice co-injected with 3 × 10^10^ particles of AAV1-EGFP-KASH. (**F**) Data were normalized to HTT mRNA in EGFP^+^ nuclei from mice injected with AAV1-CBE-mRosa26. **(F)** Mean survival and a (**G**) Kaplan-Meier curve of R6/2 mice injected with 3 × 10^10^ particles each of the AAV1-CBE-hHTT or AAV1-CBE-mRosa26 vectors (n = 14). (**D** and **F)** Bars represent means and error bars indicate S.E.M. *P < 0.05, **P < 0.01; (**D**) one-tailed unpaired t-test; (**F**) two-tailed unpaired t-test; (**G**) Log-rank Mantel-Cox test.

At four weeks after delivery, we conducted an immunohistochemical analysis to quantify CBE delivery to the striatum. Our analyses revealed that, within the striatum, ∼70% of the cells positive for the pan-neuronal marker NeuN were also positive for EGFP-KASH (**Figure 3C)** and that ∼60% of the cells positive for EGFP-KASH were positive for the Cas9 domain in the CBE protein, indicating that a proportion of cells were transduced by multiple vectors (**Figure S5**).

We next used FACS to isolate EGFP^+^ nuclei from striatal tissue of R6/2 mice injected with all three vectors, observing that ∼16% of cells from dissociated tissue were positive for EGFP (**Figure S6**). Using qPCR, we measured HTT gene expression in these enriched cell populations, finding that EGFP-KASH^+^ cells from mice injected with AAV1-CBE-hHTT had ∼50% less HTT mRNA compared to cells from control animals (P < 0.001) (**Figure 3D**), supporting the capacity for CBE-3 to lower HTT *in vivo*. We then measured by deep sequencing the frequency of C > T editing in the NF-κB binding site in a fragment of the human HTT promoter amplified from the enriched cell populations. Surprisingly, however, unlike from our cell culture studies, which revealed editing rates generally consistent with the measured decreases in HTT mRNA, we measured that only ∼2% and ∼3.5% of the analyzed reads in mice injected with AAV1-CBE-hHTT carried the -142C > T and -141C > T edits, respectively (P < 0.05; **Figure 3E**). These results thus raise the possibility that, in addition to the effects resulting from the editing of the NF-κB binding site, CBE-3 may be lowering HTT expression *in vivo* by additional mechanism(s), including potentially by physically blocking the transcription of mHTT. Additional studies are needed to unravel the *in vivo* mechanism of action of this CBE.

Because we measured a decrease mHTT in the treated animals, we also determined if CBE-3 could confer a therapeutic benefit to R6/2 mice. We therefore monitored the lifespan of four-week-old R6/2 mice injected with only the CBE-encoding AAV1 vectors, finding that animals treated with AAV1-CBE-hHTT displayed a ∼10% increase in survival compared to the mice injected with AAV1-CBE-mRosa26 (P < 0.05; **Figure 3F** and **G**). An immunohistochemical analysis of mHTT immunoreactive inclusions from the striatum of these injected animals further showed that treated mice had **∼**15% fewer CBE^+^ cells with visible mHTT inclusions compared to control mice **(Figure S7**). Thus, our results altogether demonstrate that base editors engineered to target the human HTT promoter can be delivered to an HD mouse model via dual AAV vectors and that their action can reduce HTT gene expression in transduced cells, which we show can lead to a therapeutic benefit.

### Base editing *cis*-acting sequences in the APP promoter region can perturb its expression

Finally, we sought to determine the applicability for base editing *cis*-acting sequences as an approach for perturbing target gene expression. To this end, we targeted the promoter region of the gene encoding the amyloid precursor protein (APP), a transmembrane protein whose cleavage results in the peptide fragments that comprise amyloid β (Aβ),^62^ a protein contained within the amyloid plaques that are considered a hallmark of Alzheimer’s disease (AD).^63^

Rather than relying on a CRISPRi screen to identify functional non-coding regions for targeting by base editing, we used *a priori* knowledge of the human APP promoter to design three CBE systems whose editing windows overlapped with: (i) a single conserved cytosine within a predicted binding site for the TF HSF1 (CBE-1) or (ii) multiple cytosines within GGGCGC boxes spanning -333 to -255 bps from the translation start codon (CBE-2 and -3), which have been shown to be important for modulating APP expression (**Figure 4A**).^64-67^

**Figure 4.**
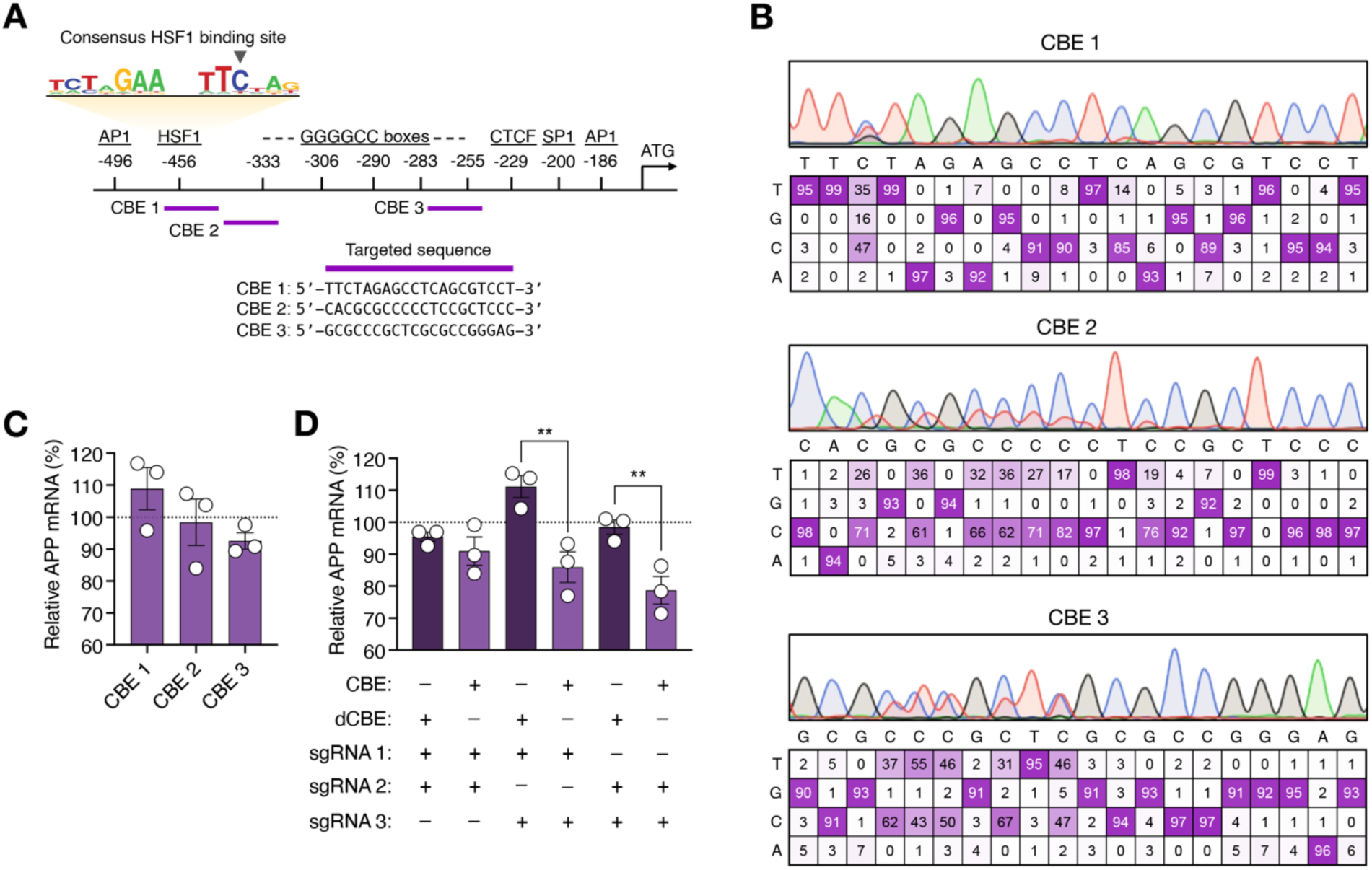
Base editing of the human APP promoter can reduce the expression of the APP gene. **(A)** (Top) Cartoon showing the positions of the binding sites for the TFs AP1, SP1, CTCF, GSF1 and AP1 and GGGCGC boxes. CBE binding sites are indicated by purple arrows. The HSF1 consensus motif is shown in the logo cartoon^85^, with black arrowheads indicating the target cytosine for CBE-1. (Bottom) Sequences targeted by the CBEs. **(B)** Sanger sequencing traces showing the editing frequencies for the candidate CBEs in HEK293T cells, as determined by EditR. **(C)** Relative APP mRNA in HEK293T cells seven days after transfection with the CBEs. **(D)** Relative APP mRNA in HEK293T cells seven days after transfection with combinations of CBEs or dCBEs. (**C** and **D**) Data were normalized to the relative APP mRNA in cells transfected with a non-targeted CBE (n = 3). Bars represent means and error bars indicate S.E.M. **P < 0.01; one-tailed unpaired t-test.

Following their transfection into HEK293T cells, we validated the editing capabilities of the CBEs using EditR,^68^ which revealed that each CBE could edit its target bases with efficiencies exceeding 30% (**Figure 4B**). Specifically, in the case of CBE-2 and CBE-3, which target GGGCGC boxes, we found that each editor could effectively mutagenize the multiple cytosines contained within their editing windows (**Figure 4B**). We next determined if the CBEs could perturb APP gene expression in HEK293T cells. Surprisingly, despite editing DNA at rates on par with the CBE targeting the NF-κB binding site in the human HTT promoter and the demonstrated importance of these elements for regulating APP,^64-66^ our qPCR analyses revealed no statistically significant differences (P > 0.05) in APP expression in HEK293T cells transfected with any of the three CBEs compared to a non-targeted CBE (**Figure 4C**). However, because TFs often operate synergistically to regulate gene expression, we reasoned that combinatorial targeting of the APP promoter could result in greater repression.^69,70^ We thus transfected HEK293T cells with combinations of the CBEs. In contrast to our earlier findings, we found that two of the three combinations (CBE-1 + CBE-3 and CBE-2 + CBE-3) significantly decreased APP mRNA in bulk-transfected cells (P < 0.05 for both; **Figure 4D**). To determine if this decrease was attributable to base-editing and not interference from the binding of the CBEs, we measured APP expression in cells transfected with dCBEs targeted to the same sites. Critically, no significant differences in APP expression were observed in cells transfected with any of the dCBE systems (P > 0.05 for all; **Figure 4D)**, indicating that the decrease in APP mRNA was attributable to the editing of the APP promoter.

In sum, our results collectively establish that base editing of *cis*-regulatory elements can facilitate target gene perturbations.

## DISCUSSION

Strategies capable of perturbing the expression of target gene(s) have the potential to enable insights into the function of genetic elements and to advance gene therapy. Here we demonstrate that CRISPR base editors,^33-35^ a programmable technology capable of introducing targeted single-base substitutions in DNA, can be harnessed to perturb target gene expression via the mutagenesis of *cis*-acting sequences in the promoter regions of genes.

We demonstrate this concept against two targets: HTT, a gene whose mutation is causative for HD and APP, a gene whose protein products comprise the plaques that are a hallmark of AD. For HTT, we show that targeted base editing of two cytosine bases in a predicted binding site for the transcription factor NF-κB decreased HTT expression. We demonstrate this effect in several contexts, including in FACS-enriched base-edited cell populations, in bulk transfected cells, and in clonally derived base-edited cell lines. For the goal of demonstrating the applicability of this concept, we also show that combinatorially targeting multiple *cis*-acting sequences in the promoter region of the APP gene, specifically the binding site for the transcription factor HSF1 and GGGCGC boxes, decreased APP gene expression. Importantly, targeting was achieved via two different approaches: in the case of HTT, we conducted a CRISPRi screen to identify an important 70-bp region in the promoter region for the HTT gene that we then subsequently tiled with CBEs, while, for APP, we used *a priori* knowledge of the human APP promoter to target annotated functional elements.^64-67^ Our results demonstrate that both methods are suitable for designing base editors to repress target gene expression.

Unlike Cas9-based methods for disrupting functional elements, base editing offers a means for mutagenizing sequences in the absence of NHEJ,^34^ a random DNA repair pathway that can generate products of varying size, which can pose risk for destroying entire TF binding sites.^15,16^ The results from our clonal analysis, in particular, demonstrate the near single nucleotide resolution of this approach, as we found that clones harboring as few as two base substitutions in the NF-κB binding site (and which also harbored no detectable CBE protein) had a two-fold decrease in target gene expression according to western blot and qPCR. Our results also highlight the specificity of this approach. Using RNA-seq, we found that only four non-target genes were differentially expressed in bulk transfected cells. Thus, despite the potential for homology with other promoter elements, this approach resulted in virtually no off-target effects in the transcriptome.

As CBE binding to a promoter region can potentially interfere with transcriptional initiation, we verified using multiple methods that the target gene perturbations observed in our cell culture studies were attributable to base-editing. In addition to a clonal analysis, which revealed no detectable CBE protein but reduced HTT mRNA and protein in the edited cell lines, we conducted studies using a catalytically inactive form of the CBE protein that still retained its ability to bind DNA (and thus interfere with transcription). Importantly, we found that only catalytically active forms of the CBEs could decrease target gene expression, as cells transfected with the deactivated equivalents of our CBE systems showed no significant differences in HTT or APP expression.

Given its toxic gain-of-function, modalities capable of lowering the mHTT protein hold promise for treating HD ^71^. Among the strategies capable of this are antisense oligonucleotides (ASOs), which have been used to lower mutant and wild-type HTT protein.^47,72-74^ However, a clinical trial determining the effectiveness of an intrathecally administered non–allele-specific ASO for HTT was halted in 2021, raising questions about the potential safety for collaterally targeting the wild-type HTT protein and whether there may exist a specific threshold for safely lowering it, particularly for intrathecally administered HTT-targeting agents.

To this end, we reasoned that base editing of the promoter region of the human HTT gene might prove an effective means for lowering wild-type and mutant HTT to a potentially safe threshold. As a first step in testing this hypothesis, we evaluated the targeted capabilities of CBE-3, the base editor engineered to target the NF-κB binding site in the human HTT promoter region, in the R6/2 mouse model of HD. While we observed in transduced nuclei a ∼50% decrease in HTT mRNA, we measured in these same cell populations editing frequencies of only ∼2% and ∼3.5% at the two targeted cytosines in the NF-κB binding site of the human HTT promoter. These results thus raise the possibility that the CBE lowered HTT *in vivo* through an additional mechanism(s) beyond DNA editing, which is at odds with the results from our cell culture experiments. Notably however, while our cell culture studies revealed editing rates generally consistent with the measured decreases in target mRNA, these experiments were conducted at time points (7 days post transfection) that may have diluted the CBE-encoding plasmids to some extent. In the case of the clonal analyses, the CBE-encoding plasmids were likely largely lost from the cell divisions, as our western blot results revealed no detectable CBE protein. By contrast, the CBE system that we delivered to mice was expected to be continuously expressed, as AAV episomes can be persistently maintained in non-dividing cells such as neurons.^75^ Thus, given its likely continuous expression, we hypothesize that the CBE protein may be preferentially physically blocking the transcription of the mHTT gene in a manner not observed in our cell culture studies. This notwithstanding, the CBE protein delivered to R6/2 mice nonetheless significantly lowered mHTT expression and provided a therapeutic benefit, as we observed that treated mice had a ∼10% increase in lifespan. However, additional studies are needed to determine the *in vivo* mechanism of action for this CBE. Finally, as demonstrated by our studies for the APP gene, we show that combinatorial targeting of multiple *cis*-acting sequences could be used to perturb expression in situations where editing of a single element had no effect on the expression of the target gene. Specifically, we reasoned that combinatorial targeting could result in a synergistic effect that could result in greater repression.^22,69^ Indeed, we found that co-transfecting CBEs targeting the HSF1 binding site and the GGGCGC boxes in the APP promoter could more effectively perturb APP expression. Along these lines, the future implementation of base editor proteins with expanded functional capabilities, ^40,41,76-78^ could provide even greater control over a target perturbation. For example, base editor variants with more restrictive editing windows could enable even greater control over the editing of a target base,^79,80^ which could lead to more effective tuning of target gene expression.

In conclusion, we establish that base editing of *cis*-regulatory elements can enable target gene perturbations. Our proof-of-concept study could pave the way for using base-editing technology to fine-tune the expression of a therapeutic gene of interest to safe and effective thresholds or to functionally interrogate non-coding elements at single base resolutions.

## MATERIALS AND METHODS

### Plasmid construction

pCMV-BE3 (Addgene #73021),^34^ pEF-BFP (Addgene #138272),^42^ pSP-gRNA (Addgene #47108),^22^ and pcDNA-dCas9,^22^ and PX552 (Addgene # 60958),^60^ were gifts from David Liu, Xiao Wang, Charles Gersbach, and Feng Zhang, respectively.

To generate the pHTT-RLuc, the target region of the human HTT promoter was PCR amplified from genomic DNA of HEK293T cells using the primers Gibson-HTT-Luc-Fwd and Gibson-HTT-Luc-Rev (**Supplementary Table S1**) and inserted between the KpnI and NheI restriction sites of psiCHECK-2 (Promega) by Gibson Assembly using the Gibson Assembly Master Mix (New England Biolabs, NEB) per the manufacturer’s instructions.

To construct the deactivated CBE expression vector, site-directed mutagenesis was performed on pCMV-BE3 by amplifying the plasmid using the oligonucleotides SDM-dCBE-Fwd and SDM-dCBE-Rev (**Supplementary Table S1**) using Phusion High-Fidelity DNA Polymerase (NEB). The resulting plasmids were incubated with DpnI (NEB) for 1 hr at 37°C and transformed into 5-alpha competent *E. coli* (NEB).

To construct the AAV1-CAG-EGFP-KASH plasmid, the CAG promoter sequence was amplified from pAAV-CAG-N-CBE-Int-U6-sgRNA,^55^ by PCR using the oligonucleotides PX552-CAG-Fwd and PX552-CAG-Rev (**Supplementary Table S1**). This amplicon was then ligated into the MluI and BamHI restriction sites of PX552.

To construct the sgRNA expression vectors, custom synthesized oligonucleotides encoding the sgRNA spacer sequences (IDT) (**Supplementary Table S1**) were phosphorylated by T4 polynucleotide kinase (NEB) for 30 min at 37°C and subsequently annealed in a thermocycler (5 min at 95°; cooled to 4°C at a rate of -0.1°C/s). Annealed oligonucleotides were then ligated into the BbsI restriction sites of pSP-gRNA or our previously described split-intein CBE-containing plasmids ^55^ using T4 DNA ligase (NEB).

The construction of all plasmids was verified by Sanger sequencing (ACGT).

### Cell culture and transfections

HEK293T cells were cultured in Dulbecco’s modified Eagle’s medium (DMEM; Corning) supplemented with 10% (v/v) fetal bovine serum (FBS; Gibco) and 1% (v/v) antibiotic-antimycotic (Gibco) in a humidified 5% CO_2_ incubator at 37°C. For transfections, HEK293T cells were seeded onto a 24-well plate at a density of 2 × 10^5^ cells per well or a 96-well plate at a density of 2 × 10^4^ cells per well. All transfections were conducted 16 hr after seeding.

For the CRISPRi screen, cells that were seeded onto a 96-well plate were transfected with 50 ng of pcDNA-dCas9, 75 ng of pSP-gRNA and 10 ng of the reporter plasmid using polyethylenimine (1 mg/mL; PEI), as previously described.^81^ For studies involving base editing of the HTT or APP promoter, cells seeded onto a 24-well plate were transfected with 750 ng of pCMV-BE3 and 500 ng of pSP-gRNA using PEI. For transient enrichment, cells seeded onto a 24-well plate were transfected with 600 ng of pCMV-BE3, 200 ng of pEF-BFP, 200 ng of pSP-gRNA and 200 ng of pSP-BFP-gRNA using PEI.

### Luciferase assays

At 72 hr after transfection, cells were lysed with Passive Lysis Buffer (Promega) and luciferase expression was determined using the Dual-Glo Luciferase Assay System (Promega) using a Synergy HTX Multimode Plate Reader (BioTek). The Renilla luminescence values in each sample was normalized to the firefly luminescence values from the same well.

### qPCR

RNA from transfected cells or striatal tissue was extracted using the PureLink RNA Mini Kit (Invitrogen) and converted to complementary DNA (cDNA) using the iScript cDNA Synthesis Kit (Bio-Rad) per the manufacturers’ instructions. qPCR was conducted in a 96-well plate using 50 ng of cDNA, 0.2 μM each of the forward and reverse primers (**Supplementary Table S1**) and iTaq Universal SYBR Green Supermix (Bio-Rad). All biological replicates were measured in technical duplicates.

### Western blot

Harvested cells were lysed by radioimmunoprecipitation assay (RIPA) buffer [0.2% IGEPAL CA-620, 0.02% SDS with Protease Inhibitor Cocktail (VWR)] and protein concentration was determined using the DC Protein Assay Kit (Bio-Rad), per the manufacturer’s instructions. 15 μg of protein per sample was then subjected to SDS-PAGE and electrophoretically transferred onto a polyvinylidene fluoride (PVDF) membrane in transfer buffer [20 mM Tris-HCl, 150 mM glycine, and 20% (v/v) methanol] for 1.5 hours at 100 V. Following the transfer, membranes were blocked using 5% (v/v) blotting-grade blocker (Bio-Rad) in Tris-Buffered Saline (10 mM Tris-HCl, 150 mM NaCl, and 0.1%, pH; TBS) with 0.05% Tween 20 (TBS-T) for 1 hr followed by an overnight incubation at 4°C with the primary antibody in blocking solution. The following primary antibodies were used: rabbit anti-HTT [EPR5526] (1:1000; Abcam, ab109115) and rabbit anti–β-actin (1:1000; Cell Signaling Technology, 4970S).

After incubation, membranes were washed with three times with TBS-T for 15 min and then incubated with goat anti-rabbit horseradish peroxidase conjugate (1:4000; ThermoFisher Scientific, 65-6120) in blocking solution for 1 hr at room temperature (RT), followed by three additional washes with TBS-T. SuperSignal West Dura Extended Duration Substrate (ThermoFisher Scientific) was then added to the membrane, with the signal visualized by automated chemiluminescence using a ChemiDoc XRS+ System (Bio-Rad). Band intensities were quantitated using Image Lab Software (Bio-Rad) and normalized to the reference band in each lane.

### Deep sequencing

DNA was extracted from harvested cells using the QuickExtract DNA Extraction Solution (Lucigen) according to the manufacturer’s instructions. Amplicon libraries were generated by PCR, subsequently indexed with Nextera adapter sequences using the KAPA2G Robust PCR Kit (Roche) and purified using the PureLink PCR Purification Kit. Libraries were then sequenced from each end by the Roy J. Carver Biotechnology Center (University of Illinois, Urbana, IL, USA) using a MiSeq Nano flow cell with a MiSeq Reagent Kit V2. The resulting FASTQ files were demultiplexed using bcl2fastq v2.17.1.14 conversion software (Illumina) per the adapter sequences. Reads were then aligned and analyzed using CRISPResso2 software. Only reads with Phred scores greater than 25 were used for analysis.

### RNA sequencing

Library construction was performed by the Roy J. Carver Biotechnology Center (University of Illinois, Urbana, IL, USA). Briefly, DNase-treated RNA was converted into barcoded polyadenylated mRNA libraries using the Kapa Hyper Stranded mRNA Sample Prep Kit (Roche). Libraries were barcoded with unique dual indexes to prevent index switching and adaptor-ligated double-stranded cDNAs were PCR-amplified for eight cycles with KAPA HiFi DNA Polymerase (Roche). Final libraries were quantitated by Qubit (ThermoFisher Scientific), and the average cDNA fragment sizes were determined on a fragment analyzer. Libraries were then diluted to 10 nM and quantitated by qPCR on a CFX Connect Real-Time qPCR system (Bio-Rad) to confirm accurate pooling of barcoded libraries and to maximize the number of clusters in the flowcell. Barcoded RNA-seq libraries were sequenced using a NovaSeq 6000 on one SP lane with single-reads 100 nt in length (Illumina). FastQ read files were generated and demultiplexed using the bcl2fastq v2.20 Conversion Software (Illumina). The quality of the demultiplexed FastQ files were then evaluated using FastQC.

Salmon version 1.5.2 was used to quasi-map reads to the transcriptome and to quantify the abundance of each transcript,^82^ using the decoy-aware method with the entire genome file as a decoy. Gene-level counts were estimated on the basis of transcript-level counts using the “lengthScaled TPM” method from the tximport package.^83^ Read counts were normalized using the trimmed mean of M-values (TMM) method from the edgeR package,^84^ and differential gene expression was tested using the limma-trend method.^84^ RNA-seq analysis was conducted by the High-Performance Biological Computing Core (HPCBio; University of Illinois, Urbana, IL, USA).

### Injections

All procedures were approved by the Illinois Institutional Animal Care and Use Committee (IACUC) at the University of Illinois and conducted in accordance with the National Institutes of Health (NIH) Guide for the Care and Use of Laboratory Animals.

Four-week-old R6/2 mice bred from male R6/2 mice [B6CBA-Tg(HDexon1)62Gpb/3J; Jackson Laboratory, Stock No: 006494] and female B6CBAF1/J mice (Jackson Laboratory, Stock No. 100011) were injected with 3 μL of AAV solution at stereotaxic coordinates anterior-posterior = 0.50 mm; medial-lateral = ±1.65 mm; and dorsal-ventral = −3.5, −3.0, and − 2.5 mm. AAV vector was manufactured and tittered by Vector Builder and injections were conducted using a drill and microinjection robot (NeuroStar). Treatment and control groups for all measurements were sex-balanced and litter-matched.

### Neuronal nuclei isolation

Neuronal nuclei were isolated from tissue as described.^57^ Briefly, harvested striatal tissue was first homogenized in 2 mL of Nuclei EZ Lysis Buffer (Sigma-Aldrich) using the KIMBLE Dounce Tissue Grinder (Sigma-Aldrich) per the manufacturer’s instructions. Following the addition of an extra 2 mL of Nuclei EZ Lysis Buffer to each homogenized tissue, samples were incubated at RT for 5 min. Homogenized tissues were then centrifuged at 500g for 5 min. After removing the supernatant, nuclei were resuspended in 4 mL of Nuclei Suspension Buffer [PBS with 100 μg/mL BSA and 3.33 μM Vibrant Dye Cycle Violet Stain (ThermoFisher Scientific)]. Nuclei were centrifuged at 500g for 1 min and resuspended in 1 mL of Nuclei Suspension Buffer for FACS.

### FACS

Harvested cells or nuclei were strained using a 35 μm filter and sorted using a BD FACSAria II Cell Sorter (Roy J. Carver Biotechnology Center Flow Cytometry Facility, University of Illinois, Urbana, IL). Cells were collected in PureLink RNA Mini Kit Lysis Buffer (Invitrogen) or DNeasy Blood & Tissue Kit Lysis Buffer (Qiagen). At least 15,000 cells or nuclei were sorted for each sample.

### Immunohistochemistry

Immunohistochemistry was performed as previously described.^55^ Harvested mouse brains were fixed in 4% paraformaldehyde (PFA) overnight at 4°C. Fixed tissues were then cut to 40 μm sagittal sections using a CM3050 S cryostat (Leica) and stored in cryoprotectant at -20°C. Prior to staining, sections were washed with PBS three times for 15 min and incubated in blocking solution [PBS with 10% (v/v) donkey serum (Abcam) and 0.5% Triton X-100] for 2 hours at RT. Sections were then stained with primary antibodies in blocking solution for 72 hr at 4°C. After incubation, sections were washed three times with PBS and incubated with secondary antibodies for 2 hr at RT. Sections were then washed three times and mounted onto slides using VECTASHIELD HardSet Antifade Mounting Medium (Vector Laboratories).

Sections were imaged using a Zeiss Observer Z1 fluorescence light microscope and a Leica TCS SP8 confocal microscope (Beckman Institute Imaging Technology Microscopy Suite, University of Illinois, Urbana, IL). Images were analyzed using ImageJ imaging software by a blinded investigator.

The following primary antibodies were used: goat anti-HA (1:250; GenScript, A00168), rabbit anti-HA (1:500; Cell Signaling Technology, 3724S), rabbit anti-NeuN (1:500; Abcam, ab177487) and mouse anti-HTT (1:50; Millipore Sigma, MAB5374).

The following secondary antibodies were used: donkey anti-rabbit Cy3 (Jackson ImmunoResearch, 711-165-152), donkey anti-goat Alexa Fluor 647 (Jackson ImmunoResearch, 705-605-147), donkey anti-mouse Alexa Fluor 488 (Jackson ImmunoResearch, 715-545-150) and donkey anti-mouse Cy3 (Jackson ImmunoResearch, 715-165-150).

### Statistical analysis

Statistical analysis was performed using GraphPad Prism 8. For *in vitro* studies, luciferase and qPCR data were analyzed by one-tailed unpaired t-test. For *in vivo* studies, survival was analyzed by a two-tailed unpaired t-test and by Kaplan-Meier analysis using the Mantel-Cox test, while qPCR and deep sequencing data were compared using a one-tailed unpaired t-test.

## DATA AVAILABILITY

Raw RNA-seq data, gene counts, and log-counts-per-million after normalization and filtering are available in the NCBI Gene Expression Omnibus.

## SUPPLEMENTARY DATA

Supplementary Data are available at NAR online.

## ACKNOWLEDGMENTS

We thank Richard Chen for helpful discussion.

## AUTHOR CONTRIBUTIONS

T.G. and P.P.P. conceived of the study; C.S.M. and C.K.W.L designed and cloned the plasmids; T.X.M. and C.S.M. conducted the luciferase assays; T.X.M., R.K. and C.K.W.L. analyzed CBE editing in HEK293T cells by deep sequencing. C.K.W.L., G.D.M and J.E.P. analyzed HTT and APP knockdown in HEK293T cells; L.V.C. analyzed RNA-seq data and performed the DEG analysis; C.K.W.L and G.D.M. bred and genotyped animals; T.X.M., C.K.W.L. and M.A.Z.C. performed stereotaxic injections; G.D.M monitored mice as a blinded investigator; C.K.W.L. and G.D.M. conducted immunohistochemistry; G.D.M. and C.K.W.L analyzed imaging data; C.K.W.L. conducted qPCR and deep sequencing measurements on striatal nuclei; T.G. and C.K.W.L. wrote the manuscript with input from all authors.

## REFERENCES

1. Buccitelli, C., and Selbach, M. (2020). mRNAs, proteins and the emerging principles of gene expression control. Nat. Rev. Genet. 21, 630–644. 10.1038/s41576-020-0258-4.

2. Wittkopp, P.J., and Kalay, G. (2012). Cis-regulatory elements: molecular mechanisms and evolutionary processes underlying divergence. Nat. Rev. Genet. 13, 59–69. 10.1038/nrg3095.

3. Cramer, P. (2019). Organization and regulation of gene transcription. Nature 573, 45–54. 10.1038/s41586-019-1517-4.

4. Yu, Z., Pandian, G.N., Hidaka, T., and Sugiyama, H. (2019). Therapeutic gene regulation using pyrrole–imidazole polyamides. Adv. Drug Deliv. Rev. 147, 66–85. https://doi.org/10.1016/j.addr.2019.02.001.

5. Matharu, N., and Ahituv, N. (2020). Modulating gene regulation to treat genetic disorders. Nat. Rev. Drug Discov. 19, 757–775. 10.1038/s41573-020-0083-7.

6. Jinek, M., Chylinski, K., Fonfara, I., Hauer, M., Doudna Jennifer, A., and Charpentier, E. (2012). A programmable dual-RNA–Guided DNA endonuclease in adaptive bacterial immunity. Science 337, 816–821. 10.1126/science.1225829.

7. Barrangou, R., Fremaux, C., Deveau, H., Richards, M., Boyaval, P., Moineau, S., Romero, D.A., and Horvath, P. (2007). CRISPR Provides Acquired Resistance Against Viruses in Prokaryotes. Science 315, 1709. 10.1126/science.1138140.

8. Wiedenheft, B., Sternberg, S.H., and Doudna, J.A. (2012). RNA-guided genetic silencing systems in bacteria and archaea. Nature 482, 331–338. 10.1038/nature10886.

9. Horvath, P., and Barrangou, R. (2010). CRISPR/Cas, the Immune System of Bacteria and Archaea. Science 327, 167–170. doi:10.1126/science.1179555.

10. Mali, P., Yang, L., Esvelt, K.M., Aach, J., Guell, M., DiCarlo, J.E., Norville, J.E., and Church, G.M. (2013). RNA-Guided Human Genome Engineering via Cas9. Science 339, 823. 10.1126/science.1232033.

11. Cong, L., Ran, F.A., Cox, D., Lin, S., Barretto, R., Habib, N., Hsu, P.D., Wu, X., Jiang, W., Marraffini, L.A., and Zhang, F. (2013). Multiplex genome engineering using CRISPR/Cas systems. Science 339, 819. 10.1126/science.1231143.

12. Cho, S.W., Kim, S., Kim, J.M., and Kim, J.-S. (2013). Targeted genome engineering in human cells with the Cas9 RNA-guided endonuclease. Nat. Biotechnol. 31, 230–232. 10.1038/nbt.2507.

13. Jiang, F., and Doudna, J.A. (2017). CRISPR–Cas9 Structures and Mechanisms. Annu. Rev. Biophys. 46, 505–529. 10.1146/annurev-biophys-062215-010822.

14. Carroll, D. (2014). Genome Engineering with Targetable Nucleases. Annu. Rev. Biochem. 83, 409–439. 10.1146/annurev-biochem-060713-035418.

15. Canver, M.C., Smith, E.C., Sher, F., Pinello, L., Sanjana, N.E., Shalem, O., Chen, D.D., Schupp, P.G., Vinjamur, D.S., Garcia, S.P., Luc, S., et al. (2015). BCL11A enhancer dissection by Cas9-mediated in situ saturating mutagenesis. Nature 527, 192–197. 10.1038/nature15521.

16. Johnston, A.D., Simões-Pires, C.A., Thompson, T.V., Suzuki, M., and Greally, J.M. (2019). Functional genetic variants can mediate their regulatory effects through alteration of transcription factor binding. Nat. Commun. 10, 3472. 10.1038/s41467-019-11412-5.

17. Fei, T., Li, W., Peng, J., Xiao, T., Chen, C.-H., Wu, A., Huang, J., Zang, C., Liu, X.S., and Brown, M. (2019). Deciphering essential cistromes using genome-wide CRISPR screens. PNAS 116, 25186–25195. doi:10.1073/pnas.1908155116.

18. Qi, Lei S., Larson, Matthew H., Gilbert, Luke A., Doudna, Jennifer A., Weissman, Jonathan S., Arkin, Adam P., and Lim, Wendell A. (2013). Repurposing CRISPR as an RNA-Guided Platform for Sequence-Specific Control of Gene Expression. Cell 152, 1173–1183. 10.1016/j.cell.2013.02.022.

19. Gilbert, Luke A., Larson, Matthew H., Morsut, L., Liu, Z., Brar, Gloria A., Torres, Sandra E., Stern-Ginossar, N., Brandman, O., Whitehead, Evan H., Doudna, Jennifer A., Lim, Wendell A., et al. (2013). CRISPR-Mediated Modular RNA-Guided Regulation of Transcription in Eukaryotes. Cell 154, 442–451. https://doi.org/10.1016/j.cell.2013.06.044.

20. Bikard, D., Jiang, W., Samai, P., Hochschild, A., Zhang, F., and Marraffini, L.A. (2013). Programmable repression and activation of bacterial gene expression using an engineered CRISPR-Cas system. Nucleic Acids Res. 41, 7429–7437. 10.1093/nar/gkt520.

21. Yeo, N.C., Chavez, A., Lance-Byrne, A., Chan, Y., Menn, D., Milanova, D., Kuo, C.-C., Guo, X., Sharma, S., Tung, A., Cecchi, R.J., et al. (2018). An enhanced CRISPR repressor for targeted mammalian gene regulation. Nat. Methods 15, 611–616. 10.1038/s41592-018-0048-5.

22. Perez-Pinera, P., Kocak, D.D., Vockley, C.M., Adler, A.F., Kabadi, A.M., Polstein, L.R., Thakore, P.I., Glass, K.A., Ousterout, D.G., Leong, K.W., Guilak, F., et al. (2013). RNA-guided gene activation by CRISPR-Cas9–based transcription factors. Nat. Methods 10, 973–976. 10.1038/nmeth.2600.

23. Maeder, M.L., Linder, S.J., Cascio, V.M., Fu, Y., Ho, Q.H., and Joung, J.K. (2013). CRISPR RNA–guided activation of endogenous human genes. Nat. Methods 10, 977–979. 10.1038/nmeth.2598.

24. Chavez, A., Scheiman, J., Vora, S., Pruitt, B.W., Tuttle, M., P R Iyer, E., Lin, S., Kiani, S., Guzman, C.D., Wiegand, D.J., Ter-Ovanesyan, D., et al. (2015). Highly efficient Cas9-mediated transcriptional programming. Nat. Methods 12, 326–328. 10.1038/nmeth.3312.

25. Konermann, S., Brigham, M.D., Trevino, A.E., Joung, J., Abudayyeh, O.O., Barcena, C., Hsu, P.D., Habib, N., Gootenberg, J.S., Nishimasu, H., Nureki, O., et al. (2015). Genome-scale transcriptional activation by an engineered CRISPR-Cas9 complex. Nature 517, 583–588. 10.1038/nature14136.

26. Tanenbaum, Marvin E., Gilbert, Luke A., Qi, Lei S., Weissman, Jonathan S., and Vale, Ronald D. (2014). A Protein-Tagging System for Signal Amplification in Gene Expression and Fluorescence Imaging. Cell 159, 635–646. https://doi.org/10.1016/j.cell.2014.09.039.

27. van Overbeek, M., Capurso, D., Carter, Matthew M., Thompson, Matthew S., Frias, E., Russ, C., Reece-Hoyes, John S., Nye, C., Gradia, S., Vidal, B., Zheng, J., et al. (2016). DNA Repair Profiling Reveals Nonrandom Outcomes at Cas9-Mediated Breaks. Mol. Cell 63, 633–646. https://doi.org/10.1016/j.molcel.2016.06.037.

28. Allen, F., Crepaldi, L., Alsinet, C., Strong, A.J., Kleshchevnikov, V., De Angeli, P., Páleníková, P., Khodak, A., Kiselev, V., Kosicki, M., Bassett, A.R., et al. (2019). Predicting the mutations generated by repair of Cas9-induced double-strand breaks. Nat. Biotechnol. 37, 64–72. 10.1038/nbt.4317.

29. Chakrabarti, A.M., Henser-Brownhill, T., Monserrat, J., Poetsch, A.R., Luscombe, N.M., and Scaffidi, P. (2019). Target-Specific Precision of CRISPR-Mediated Genome Editing. Mol. Cell 73, 699-713.e696. 10.1016/j.molcel.2018.11.031.

30. Saleh-Gohari, N., and Helleday, T. (2004). Conservative homologous recombination preferentially repairs DNA double-strand breaks in the S phase of the cell cycle in human cells. Nucleic Acids Res. 32, 3683–3688. 10.1093/nar/gkh703.

31. Miyaoka, Y., Berman, J.R., Cooper, S.B., Mayerl, S.J., Chan, A.H., Zhang, B., Karlin-Neumann, G.A., and Conklin, B.R. (2016). Systematic quantification of HDR and NHEJ reveals effects of locus, nuclease, and cell type on genome-editing. Sci. Rep. 6, 23549. 10.1038/srep23549.

32. Fu, Y.-W., Dai, X.-Y., Wang, W.-T., Yang, Z.-X., Zhao, J.-J., Zhang, J.-P., Wen, W., Zhang, F., Oberg, K.C., Zhang, L., Cheng, T., et al. (2021). Dynamics and competition of CRISPR–Cas9 ribonucleoproteins and AAV donor-mediated NHEJ, MMEJ and HDR editing. Nucleic Acids Res. 49, 969–985. 10.1093/nar/gkaa1251.

33. Rees, H.A., and Liu, D.R. (2018). Base editing: precision chemistry on the genome and transcriptome of living cells. Nat. Rev. Genet. 19, 770–788. 10.1038/s41576-018-0059-1.

34. Komor, A.C., Kim, Y.B., Packer, M.S., Zuris, J.A., and Liu, D.R. (2016). Programmable editing of a target base in genomic DNA without double-stranded DNA cleavage. Nature 533, 420–424. 10.1038/nature17946.

35. Gaudelli, N.M., Komor, A.C., Rees, H.A., Packer, M.S., Badran, A.H., Bryson, D.I., and Liu, D.R. (2017). Programmable base editing of A•T to G•C in genomic DNA without DNA cleavage. Nature 551, 464–471. 10.1038/nature24644.

36. Tabrizi, S.J., Flower, M.D., Ross, C.A., and Wild, E.J. (2020). Huntington disease: new insights into molecular pathogenesis and therapeutic opportunities. Nat. Rev. Neurol. 16, 529–546. 10.1038/s41582-020-0389-4.

37. Holzmann, C., Schmidt, T., Thiel, G., Epplen, J.T., and Riess, O. (2001). Functional characterization of the human Huntington’s disease gene promoter. Mol. Brain Res. 92, 85–97. https://doi.org/10.1016/S0169-328X(01)00149-8.

38. Becanovic, K., Nørremølle, A., Neal, S.J., Kay, C., Collins, J.A., Arenillas, D., Lilja, T., Gaudenzi, G., Manoharan, S., Doty, C.N., Beck, J., et al. (2015). A SNP in the HTT promoter alters NF-κB binding and is a bidirectional genetic modifier of Huntington disease. Nat. Neurosci. 18, 807–816. 10.1038/nn.4014.

39. Khoshnan, A., Ko, J., Watkin, E.E., Paige, L.A., Reinhart, P.H., and Patterson, P.H. (2004). Activation of the IκB Kinase Complex and Nuclear Factor-κB Contributes to Mutant Huntingtin Neurotoxicity. J. Neurosci. 24, 7999. 10.1523/JNEUROSCI.2675-04.2004.

40. Kim, Y.B., Komor, A.C., Levy, J.M., Packer, M.S., Zhao, K.T., and Liu, D.R. (2017). Increasing the genome-targeting scope and precision of base editing with engineered Cas9-cytidine deaminase fusions. Nat. Biotechnol. 35, 371–376. 10.1038/nbt.3803.

41. Komor, A.C., Zhao, K.T., Packer, M.S., Gaudelli, N.M., Waterbury, A.L., Koblan, L.W., Kim, Y.B., Badran, A.H., and Liu, D.R. (2017). Improved base excision repair inhibition and bacteriophage Mu Gam protein yields C:G-to-T:A base editors with higher efficiency and product purity. Sci. Adv. 3, eaao4774. doi:10.1126/sciadv.aao4774.

42. Standage-Beier, K., Tekel, S.J., Brookhouser, N., Schwarz, G., Nguyen, T., Wang, X., and Brafman, D.A. (2019). A transient reporter for editing enrichment (TREE) in human cells. Nucleic Acids Res. 47, e120–e120. 10.1093/nar/gkz713.

43. Coelho, M.A., Li, S., Pane, L.S., Firth, M., Ciotta, G., Wrigley, J.D., Cuomo, M.E., Maresca, M., and Taylor, B.J.M. (2018). BE-FLARE: a fluorescent reporter of base editing activity reveals editing characteristics of APOBEC3A and APOBEC3B. BMC Biol. 16, 150. 10.1186/s12915-018-0617-1.

44. Zuo, E., Sun, Y., Yuan, T., He, B., Zhou, C., Ying, W., Liu, J., Wei, W., Zeng, R., Li, Y., and Yang, H. (2020). A rationally engineered cytosine base editor retains high on-target activity while reducing both DNA and RNA off-target effects. Nat. Methods 17, 600–604. 10.1038/s41592-020-0832-x.

45. Lambert, S.A., Jolma, A., Campitelli, L.F., Das, P.K., Yin, Y., Albu, M., Chen, X., Taipale, J., Hughes, T.R., and Weirauch, M.T. (2018). The Human Transcription Factors. Cell 172, 650–665. https://doi.org/10.1016/j.cell.2018.01.029.

46. Mangiarini, L., Sathasivam, K., Seller, M., Cozens, B., Harper, A., Hetherington, C., Lawton, M., Trottier, Y., Lehrach, H., Davies, S.W., and Bates, G.P. (1996). Exon 1 of the HD gene with an expanded CAG repeat is sufficient to cause a progressive neurological phenotype in transgenic mice. Cell 87, 493–506. 10.1016/s0092-8674(00)81369-0.

47. Kordasiewicz, H.B., Stanek, L.M., Wancewicz, E.V., Mazur, C., McAlonis, M.M., Pytel, K.A., Artates, J.W., Weiss, A., Cheng, S.H., Shihabuddin, L.S., Hung, G., et al. (2012). Sustained therapeutic reversal of Huntington’s disease by transient repression of huntingtin synthesis. Neuron 74, 1031–1044. 10.1016/j.neuron.2012.05.009.

48. Davies, S.W., Turmaine, M., Cozens, B.A., DiFiglia, M., Sharp, A.H., Ross, C.A., Scherzinger, E., Wanker, E.E., Mangiarini, L., and Bates, G.P. (1997). Formation of Neuronal Intranuclear Inclusions Underlies the Neurological Dysfunction in Mice Transgenic for the HD Mutation. Cell 90, 537–548. 10.1016/S0092-8674(00)80513-9.

49. Harper, S.Q., Staber, P.D., He, X., Eliason, S.L., Martins, I.H., Mao, Q., Yang, L., Kotin, R.M., Paulson, H.L., and Davidson, B.L. (2005). RNA interference improves motor and neuropathological abnormalities in a Huntington’s disease mouse model. PNAS 102, 5820–5825. 10.1073/pnas.0501507102.

50. Ekman, F.K., Ojala, D.S., Adil, M.M., Lopez, P.A., Schaffer, D.V., and Gaj, T. (2019). CRISPR-Cas9-Mediated Genome Editing Increases Lifespan and Improves Motor Deficits in a Huntington’s Disease Mouse Model. Mol. Ther. Nucleic Acids 17, 829–839. 10.1016/j.omtn.2019.07.009.

51. Boudreau, R.L., McBride, J.L., Martins, I., Shen, S., Xing, Y., Carter, B.J., and Davidson, B.L. (2009). Nonallele-specific silencing of mutant and wild-type huntingtin demonstrates therapeutic efficacy in Huntington’s disease mice. Mol. Ther. 17, 1053–1063. 10.1038/mt.2009.17.

52. McBride, J.L., Boudreau, R.L., Harper, S.Q., Staber, P.D., Monteys, A.M., Martins, I., Gilmore, B.L., Burstein, H., Peluso, R.W., Polisky, B., Carter, B.J., et al. (2008). Artificial miRNAs mitigate shRNA-mediated toxicity in the brain: Implications for the therapeutic development of RNAi. PNAS 105, 5868. 10.1073/pnas.0801775105.

53. Nieuwenhuis, B., Haenzi, B., Hilton, S., Carnicer-Lombarte, A., Hobo, B., Verhaagen, J., and Fawcett, J.W. (2021). Optimization of adeno-associated viral vector-mediated transduction of the corticospinal tract: comparison of four promoters. Gene Ther. 28, 56–74. 10.1038/s41434-020-0169-1.

54. Powell, J.E., Lim, C.K.W., Krishnan, R., McCallister, T.X., Saporito-Magriña, C., Zeballos, M.A., McPheron, G.D., and Gaj, T. (2022). Targeted gene silencing in the nervous system with CRISPR-Cas13. Sci. Adv. 8, eabk2485. doi:10.1126/sciadv.abk2485.

55. Lim, C.K.W., Gapinske, M., Brooks, A.K., Woods, W.S., Powell, J.E., Zeballos C, M.A., Winter, J., Perez-Pinera, P., and Gaj, T. (2020). Treatment of a mouse model of ALS by in vivo base editing. Mol. Ther. 28, 1177–1189. https://doi.org/10.1016/j.ymthe.2020.01.005.

56. Winter, J., Luu, A., Gapinske, M., Manandhar, S., Shirguppe, S., Woods, W.S., Song, J.S., and Perez-Pinera, P. (2019). Targeted exon skipping with AAV-mediated split adenine base editors. Cell Discov. 5, 41. 10.1038/s41421-019-0109-7.

57. Levy, J.M., Yeh, W.-H., Pendse, N., Davis, J.R., Hennessey, E., Butcher, R., Koblan, L.W., Comander, J., Liu, Q., and Liu, D.R. (2020). Cytosine and adenine base editing of the brain, liver, retina, heart and skeletal muscle of mice via adeno-associated viruses. Nat. Biomed. Eng. 4, 97–110. 10.1038/s41551-019-0501-5.

58. Villiger, L., Grisch-Chan, H.M., Lindsay, H., Ringnalda, F., Pogliano, C.B., Allegri, G., Fingerhut, R., Häberle, J., Matos, J., Robinson, M.D., Thöny, B., et al. (2018). Treatment of a metabolic liver disease by in vivo genome base editing in adult mice. Nat. Med. 24, 1519–1525. 10.1038/s41591-018-0209-1.

59. Chen, Y., Zhi, S., Liu, W., Wen, J., Hu, S., Cao, T., Sun, H., Li, Y., Huang, L., Liu, Y., Liang, P., et al. (2020). Development of Highly Efficient Dual-AAV Split Adenosine Base Editor for In Vivo Gene Therapy. Small Methods 4, 2000309. https://doi.org/10.1002/smtd.202000309.

60. Swiech, L., Heidenreich, M., Banerjee, A., Habib, N., Li, Y., Trombetta, J., Sur, M., and Zhang, F. (2015). In vivo interrogation of gene function in the mammalian brain using CRISPR-Cas9. Nat. Biotechnol. 33, 102–106. 10.1038/nbt.3055.

61. Östlund, C., Folker, E.S., Choi, J.C., Gomes, E.R., Gundersen, G.G., and Worman, H.J. (2009). Dynamics and molecular interactions of linker of nucleoskeleton and cytoskeleton (LINC) complex proteins. J. Cell Sci. 122, 4099–4108. 10.1242/jcs.057075.

62. Haass, C., and Selkoe, D.J. (1993). Cellular processing of β-amyloid precursor protein and the genesis of amyloid β-peptide. Cell 75, 1039–1042. https://doi.org/10.1016/0092-8674(93)90312-E.

63. Querfurth, H.W., and LaFerla, F.M. (2010). Alzheimer’s Disease. N. Engl. J. Med. 362, 329–344. 10.1056/NEJMra0909142.

64. Lahiri, D.K., and Robakis, N.K. (1991). The promoter activity of the gene encoding Alzheimer β-amyloid precursor protein (APP) is regulated by two blocks of upstream sequences. Mol. Brain Res. 9, 253–257. https://doi.org/10.1016/0169-328X(91)90009-M.

65. Pollwein, P., Masters, C.L., and Beyreuther, K. (1992). The expression of the amyloid precursor protein (APP) is regulated by two GC-elements in the promoter. Nucleic Acids Res. 20, 63–68. 10.1093/nar/20.1.63.

66. Salbaum, J.M., Weidemann, A., Lemaire, H.G., Masters, C.L., and Beyreuther, K. (1988). The promoter of Alzheimer’s disease amyloid A4 precursor gene. EMBO J. 7, 2807–2813. https://doi.org/10.1002/j.1460-2075.1988.tb03136.x.

67. La Fauci, G., Lahiri, D.K., Salton, S.R.J., and Robakis, N.K. (1989). Characterization of the 5′-end region and the first two exons of the β-protein precursor gene. Biochem. Biophys. Res. Commun. 159, 297–304. https://doi.org/10.1016/0006-291X(89)92437-6.

68. Kluesner, M.G., Nedveck, D.A., Lahr, W.S., Garbe, J.R., Abrahante, J.E., Webber, B.R., and Moriarity, B.S. (2018). EditR: A method to quantify base editing from Sanger sequencing. CRISPR J. 1, 239–250. 10.1089/crispr.2018.0014.

69. Carey, M. (1998). The Enhanceosome and Transcriptional Synergy. Cell 92, 5–8. 10.1016/S0092-8674(00)80893-4.

70. Cumbo, F., Vergni, D., and Santoni, D. (2017). Investigating transcription factor synergism in humans. DNA Res. 25, 103–112. 10.1093/dnares/dsx041.

71. Tabrizi, S.J., Ghosh, R., and Leavitt, B.R. (2019). Huntingtin Lowering Strategies for Disease Modification in Huntington’s Disease. Neuron 101, 801–819. 10.1016/j.neuron.2019.01.039.

72. Tabrizi, S.J., Leavitt, B.R., Landwehrmeyer, G.B., Wild, E.J., Saft, C., Barker, R.A., Blair, N.F., Craufurd, D., Priller, J., Rickards, H., Rosser, A., et al. (2019). Targeting Huntingtin Expression in Patients with Huntington’s Disease. N. Engl. J. Med. 380, 2307–2316. 10.1056/NEJMoa1900907.

73. Stanek, L.M., Yang, W., Angus, S., Sardi, P.S., Hayden, M.R., Hung, G.H., Bennett, C.F., Cheng, S.H., and Shihabuddin, L.S. (2013). Antisense Oligonucleotide-Mediated Correction of Transcriptional Dysregulation is Correlated with Behavioral Benefits in the YAC128 Mouse Model of Huntington’s Disease. J. Huntington’s Dis. 2, 217–228. 10.3233/JHD-130057.

74. Southwell, A.L., Kordasiewicz, H.B., Langbehn, D., Skotte, N.H., Parsons, M.P., Villanueva, E.B., Caron, N.S., Østergaard, M.E., Anderson, L.M., Xie, Y., Cengio, L.D., et al. (2018). Huntingtin suppression restores cognitive function in a mouse model of Huntington’s disease. Sci. Transl. Med. 10, eaar3959. doi:10.1126/scitranslmed.aar3959.

75. Afione, S.A., Conrad, C.K., Kearns, W.G., Chunduru, S., Adams, R., Reynolds, T.C., Guggino, W.B., Cutting, G.R., Carter, B.J., and Flotte, T.R. (1996). In vivo model of adeno-associated virus vector persistence and rescue. J. Virol. 70, 3235–3241. 10.1128/JVI.70.5.3235-3241.1996.

76. Yu, Y., Leete, T.C., Born, D.A., Young, L., Barrera, L.A., Lee, S.-J., Rees, H.A., Ciaramella, G., and Gaudelli, N.M. (2020). Cytosine base editors with minimized unguided DNA and RNA off-target events and high on-target activity. Nat. Commun. 11, 2052. 10.1038/s41467-020-15887-5.

77. Koblan, L.W., Doman, J.L., Wilson, C., Levy, J.M., Tay, T., Newby, G.A., Maianti, J.P., Raguram, A., and Liu, D.R. (2018). Improving cytidine and adenine base editors by expression optimization and ancestral reconstruction. Nat. Biotechnol. 36, 843–846. 10.1038/nbt.4172.

78. Thuronyi, B.W., Koblan, L.W., Levy, J.M., Yeh, W.-H., Zheng, C., Newby, G.A., Wilson, C., Bhaumik, M., Shubina-Oleinik, O., Holt, J.R., and Liu, D.R. (2019). Continuous evolution of base editors with expanded target compatibility and improved activity. Nat. Biotechnol. 37, 1070–1079. 10.1038/s41587-019-0193-0.

79. Tan, J., Zhang, F., Karcher, D., and Bock, R. (2019). Engineering of high-precision base editors for site-specific single nucleotide replacement. Nat. Commun. 10, 439. 10.1038/s41467-018-08034-8.

80. Huang, T.P., Zhao, K.T., Miller, S.M., Gaudelli, N.M., Oakes, B.L., Fellmann, C., Savage, D.F., and Liu, D.R. (2019). Circularly permuted and PAM-modified Cas9 variants broaden the targeting scope of base editors. Nat. Biotechnol. 37, 626–631. 10.1038/s41587-019-0134-y.

81. Gaj, T., and Schaffer, D.V. (2016). Adeno-Associated Virus–Mediated Delivery of CRISPR–Cas Systems for Genome Engineering in Mammalian Cells. Cold Spring Harb. Protoc. 2016, pdb.prot086868. 10.1101/pdb.prot086868.

82. Patro, R., Duggal, G., Love, M.I., Irizarry, R.A., and Kingsford, C. (2017). Salmon provides fast and bias-aware quantification of transcript expression. Nat. Methods 14, 417–419. 10.1038/nmeth.4197.

83. Soneson, C., Love, M.I., and Robinson, M.D. (2015). Differential analyses for RNA-seq: transcript-level estimates improve gene-level inferences. F1000Res 4, 1521–1521. 10.12688/f1000research.7563.2.

84. Chen, Y., Lun, A.T.L., and Smyth, G.K. (2016). From reads to genes to pathways: differential expression analysis of RNA-Seq experiments using Rsubread and the edgeR quasi-likelihood pipeline. F1000Res 5, 1438–1438. 10.12688/f1000research.8987.2.

85. Zamdborg, L., and Ma, P. (2009). Discovery of protein–DNA interactions by penalized multivariate regression. Nucleic Acids Res. 37, 5246–5254. 10.1093/nar/gkp554.

