## Supplementary Information for "CRISPR base editing of *cis*-regulatory elements enables target gene perturbations"


Colin K.W. Lim[1], Tristan X. McCallister[1], Christian Saporito-Magriña[1], Garrett D. McPheron[1], Ramya Krishnan[1], M. Alejandra Zeballos C[1], Jackson E. Powell[1], Lindsay V. Clark[2], Pablo Perez-Pinera[1,3,4,5,*] and Thomas Gaj[1,3,*]

1 Department of Bioengineering, University of Illinois, Urbana, IL 61801, USA

2 Roy J. Carver Biotechnology Center, University of Illinois at Urbana-Champaign, Urbana, IL, 61801, USA

3 Carl R. Woese Institute for Genomic Biology, University of Illinois, Urbana, IL 61801, USA

4 Department of Biomedical and Translational Sciences, Carle-Illinois College of Medicine, University of Illinois, Urbana, IL 61801, USA

5 Cancer Center at Illinois, University of Illinois, Urbana, IL 61801, USA

* Correspondence should be addressed to:

P.P.P and T.G.

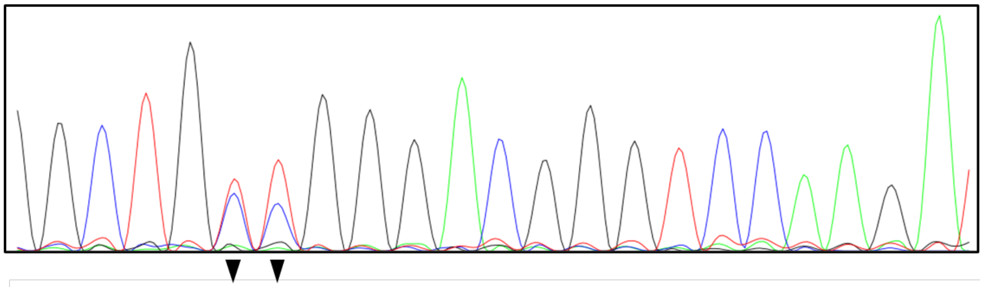

| | G | C | T | G | C | C | G | G | G | A | C | G | G | G | T | C | C | A | A | G | A |
|---|---|---|---|---|---|---|---|---|---|---|---|---|---|---|---|---|---|---|---|---|---|
| T | 6 | 9 | 90 | 4 | 51 | 61 | 4 | 3 | 4 | 3 | 9 | 8 | 5 | 8 | 90 | 11 | 9 | 10 | 6 | 9 | 3 |
| G | 87 | 4 | 5 | 90 | 5 | 6 | 92 | 91 | 87 | 2 | 4 | 83 | 89 | 85 | 2 | 2 | 2 | 5 | 6 | 76 | 4 |
| C | 5 | 83 | 4 | 3 | 41 | 32 | 2 | 2 | 3 | 2 | 83 | 6 | 3 | 3 | 5 | 83 | 83 | 4 | 0 | 4 | 0 |
| A | 2 | 4 | 1 | 2 | 4 | 2 | 2 | 4 | 6 | 93 | 4 | 4 | 3 | 4 | 3 | 5 | 7 | 81 | 88 | 12 | 93 |

**Supplementary Figure S1. Transient enrichment increased the percentage of HEK293T cells with edits within the NF-κB binding site in the HTT promoter**. A representative Sanger sequencing trace and the corresponding EditR analysis of the base editing frequencies in the NF-κB binding site in the HTT promoter from HEK293T cells transfected with CBE-3 and the transient enrichment reporter system (pEF-BFP and the BFP-targeting sgRNA). Arrowheads indicate the target cytosines in the NF-κB binding site.

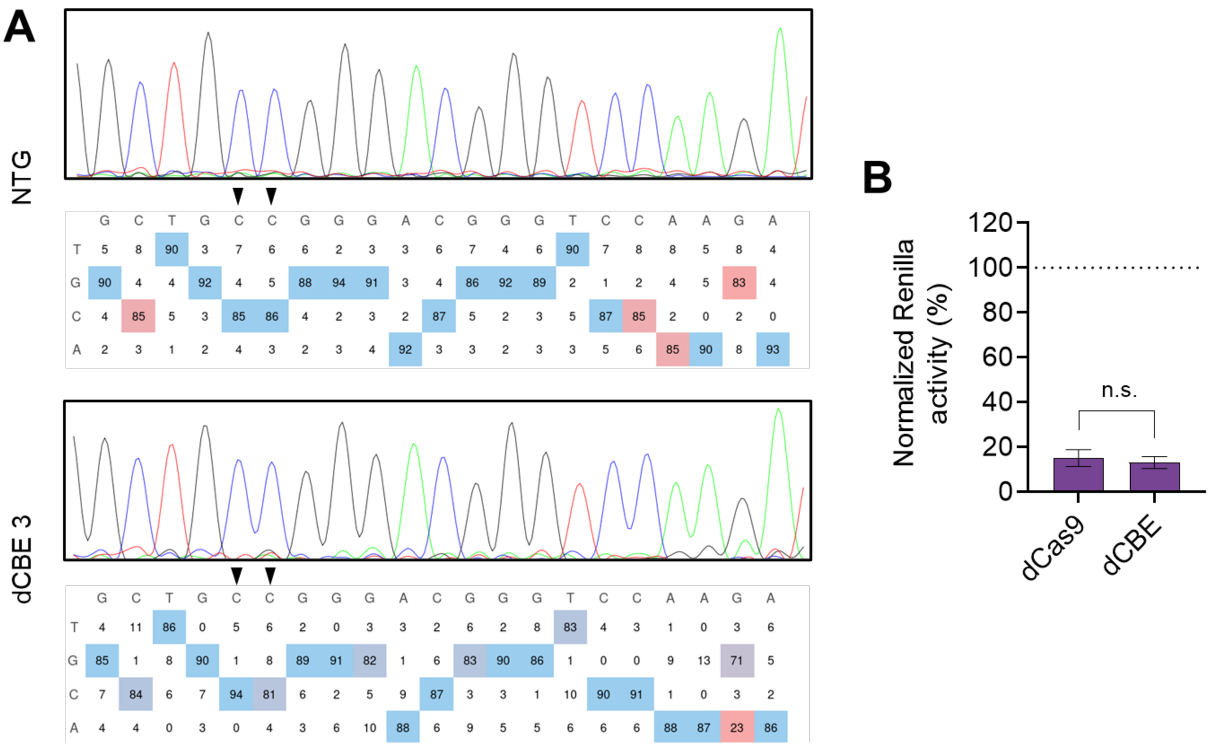


**Supplementary Figure S2**. **Deactivated CBEs (dCBEs) do not edit DNA but retain the ability to bind DNA. (A)** A representative Sanger sequencing traces and the corresponding EditR analysis of the base editing frequencies in the NF-κB binding site in the HTT promoter from HEK293T cells transfected with a (top) non-targeting CBE or (bottom) the dCBE equivalent of CBE-3 (dCBE-3). Arrowheads indicate the target cytosines in the NF-κB binding site. **(B)** Normalized Renilla luciferase expression in HEK293T cells after transfection with the pHTT-RLuc reporter plasmid and expression vectors encoding sgRNA 8 from **Figure 1B (**which overlaps with the NF-κB binding site) and dCas9 or dCBE. Renilla expression was normalized to firefly luciferase, and all relative values were normalized to cells trasfected with pHTT-RLuc and dCas9 with a non-targeted sgRNA (n = 3). Bars indicate the means and error bars indicate the S.D.

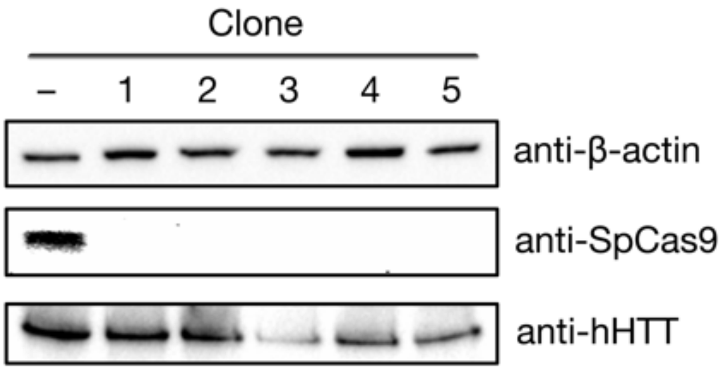


**Supplementary Figure S3. Base-edited clones expressed less HTT protein than negative control cells and had no detectable CBE protein**. Western blot of cell lysate from HEK293T clones originally transfected with CBE-3. "–" indicates HEK293T cells transfected with a non-targeted CBE, which were harvested at 72 hr post-transfection. Quantitation of the western blot is presented in **Figure 2D**.

**A**

| ID | Target sequence | Location |
|---|---|---|
| hHTT: | 5’-CTGCCGGGACGGGTCCAAGA-3’ | chr4:+3074699 |
| OT1: | 5’-CTGATGGCAAGGGTCCAAGA-3’ | chr6:-19633866 |
| OT2: | 5’-CTGTTGTGAGGGGTCCAAGA-3’ | chr1:-1227771 |
| OT3: | 5’-CTCCAGGGAGGGCTCCAAGA-3’ | chr11:-72828586 |
| OT4: | 5’-CTTCTGGGATGGATCCAAGA-3’ | chr1:-167529418 |
| OT5: | 5’-GTGCCTGGCTGGGTCCAAGA-3’ | chr20:+46302556 |

**B**

OT1

| | C | T | G | A | T | G | G | C | A | A | G | G | G | T | C | C | A | A | G | A |
|---|---|---|---|---|---|---|---|---|---|---|---|---|---|---|---|---|---|---|---|---|
| A | 0.055 | 0.210 | 0.180 | 99.164 | 0.158 | 0.291 | 0.249 | 0.014 | 99.086 | 99.195 | 0.120 | 0.240 | 0.198 | 0.157 | 0.033 | 0.053 | 99.191 | 99.217 | 0.124 | 99.252 |
| C | 99.716 | 0.507 | 0.010 | 0.034 | 0.512 | 0.004 | 0.031 | 99.661 | 0.035 | 0.028 | 0.010 | 0.016 | 0.019 | 0.639 | 99.749 | 99.574 | 0.050 | 0.036 | 0.011 | 0.018 |
| G | 0.012 | 0.044 | 99.779 | 0.586 | 0.029 | 99.672 | 99.622 | 0.019 | 0.703 | 0.556 | 99.782 | 99.647 | 99.740 | 0.061 | 0.017 | 0.015 | 0.596 | 0.557 | 99.837 | 0.607 |
| T | 0.209 | 99.238 | 0.029 | 0.215 | 99.299 | 0.033 | 0.064 | 0.293 | 0.176 | 0.159 | 0.047 | 0.097 | 0.043 | 99.134 | 0.200 | 0.216 | 0.163 | 0.174 | 0.027 | 0.122 |

OT2

| | C | T | G | T | T | G | T | G | A | G | G | G | G | T | C | C | A | A | G | A |
|---|---|---|---|---|---|---|---|---|---|---|---|---|---|---|---|---|---|---|---|---|
| A | 0.021 | 0.230 | 0.193 | 0.134 | 0.078 | 0.223 | 0.110 | 0.173 | 98.998 | 0.148 | 0.178 | 0.296 | 0.252 | 0.101 | 0.044 | 0.038 | 99.258 | 99.153 | 0.131 | 99.231 |
| C | 99.782 | 0.636 | 0.009 | 0.675 | 0.649 | 0.003 | 0.627 | 0.005 | 0.023 | 0.005 | 0.007 | 0.013 | 0.015 | 0.776 | 99.732 | 99.621 | 0.041 | 0.026 | 0.009 | 0.016 |
| G | 0.003 | 0.030 | 99.784 | 0.019 | 0.025 | 99.764 | 0.037 | 99.775 | 0.740 | 99.817 | 99.679 | 99.628 | 99.719 | 0.023 | 0.016 | 0.006 | 0.595 | 0.648 | 99.843 | 0.628 |
| T | 0.174 | 99.101 | 0.013 | 99.171 | 99.240 | 0.008 | 99.193 | 0.014 | 0.222 | 0.025 | 0.060 | 0.063 | 0.015 | 99.095 | 0.208 | 0.257 | 0.106 | 0.156 | 0.014 | 0.125 |

OT3

| | C | T | C | C | A | G | G | G | A | G | G | G | C | T | C | C | A | A | G | A |
|---|---|---|---|---|---|---|---|---|---|---|---|---|---|---|---|---|---|---|---|---|
| A | 0.035 | 0.099 | 0.037 | 0.035 | 99.378 | 0.118 | 0.132 | 0.130 | 99.203 | 0.153 | 0.219 | 0.227 | 0.032 | 0.087 | 0.051 | 0.023 | 99.407 | 99.336 | 0.118 | 99.311 |
| C | 99.759 | 0.648 | 99.803 | 99.604 | 0.035 | 0.004 | 0.010 | 0.009 | 0.031 | 0.005 | 0.006 | 0.026 | 99.801 | 0.776 | 99.779 | 99.773 | 0.029 | 0.028 | 0.003 | 0.027 |
| G | 0.007 | 0.038 | 0.010 | 0.009 | 0.497 | 99.852 | 99.831 | 99.779 | 0.602 | 99.812 | 99.630 | 99.722 | 0.006 | 0.031 | 0.013 | 0.006 | 0.499 | 0.518 | 99.860 | 0.557 |
| T | 0.190 | 99.195 | 0.149 | 0.176 | 0.086 | 0.026 | 0.026 | 0.031 | 0.139 | 0.029 | 0.055 | 0.024 | 0.161 | 99.098 | 0.156 | 0.149 | 0.064 | 0.104 | 0.017 | 0.098 |

OT4

| | C | T | T | C | T | G | G | G | A | T | G | G | A | T | C | C | A | A | G | A |
|---|---|---|---|---|---|---|---|---|---|---|---|---|---|---|---|---|---|---|---|---|
| A | 0.011 | 0.059 | 0.086 | 0.029 | 0.127 | 0.149 | 0.172 | 0.128 | 99.260 | 0.058 | 0.186 | 0.177 | 99.289 | 0.054 | 0.038 | 0.033 | 99.409 | 99.298 | 0.108 | 99.331 |
| C | 99.874 | 0.671 | 0.473 | 99.845 | 0.472 | 0.004 | 0.017 | 0.014 | 0.023 | 0.453 | 0.009 | 0.010 | 0.015 | 0.596 | 99.788 | 99.753 | 0.032 | 0.036 | 0.002 | 0.017 |
| G | 0.003 | 0.026 | 0.018 | 0.003 | 0.040 | 99.828 | 99.729 | 99.643 | 0.620 | 0.031 | 99.672 | 99.742 | 0.622 | 0.019 | 0.015 | 0.011 | 0.509 | 0.560 | 99.861 | 0.566 |
| T | 0.113 | 99.243 | 99.419 | 0.121 | 99.360 | 0.019 | 0.080 | 0.034 | 0.092 | 99.457 | 0.026 | 0.071 | 0.071 | 99.328 | 0.159 | 0.161 | 0.050 | 0.092 | 0.027 | 0.083 |

OT5

| | G | T | G | C | C | T | G | G | C | T | G | G | G | T | C | C | A | A | G | A |
|---|---|---|---|---|---|---|---|---|---|---|---|---|---|---|---|---|---|---|---|---|
| A | 0.205 | 0.185 | 0.210 | 0.048 | 0.025 | 0.195 | 0.239 | 0.219 | 0.018 | 0.100 | 0.194 | 0.265 | 0.205 | 0.092 | 0.029 | 0.033 | 99.338 | 99.292 | 0.115 | 99.288 |
| C | 0.005 | 0.682 | 0.009 | 99.688 | 99.759 | 0.542 | 0.005 | 0.015 | 99.738 | 0.435 | 0.006 | 0.018 | 0.018 | 0.708 | 99.767 | 99.666 | 0.049 | 0.030 | 0.004 | 0.029 |
| G | 99.771 | 0.037 | 99.755 | 0.044 | 0.004 | 0.039 | 99.735 | 99.651 | 0.003 | 0.043 | 99.519 | 99.625 | 99.760 | 0.033 | 0.013 | 0.005 | 0.537 | 0.543 | 99.858 | 0.587 |
| T | 0.018 | 99.090 | 0.019 | 0.220 | 0.189 | 99.214 | 0.021 | 0.028 | 0.239 | 99.420 | 0.034 | 0.089 | 0.017 | 99.159 | 0.191 | 0.230 | 0.074 | 0.116 | 0.021 | 0.096 |

**Supplementary Figure S4. CBE-3 did not edit computationally predicted off-target sites**. **(A)** DNA sequences and chromosomal locations of the target sequence for CBE-3 and five potential off-target sites identified using an algorithm described Hsu *et al* (1). **(B)** Tables showing the nucleotide frequencies of each base, obtained by deep seqeuncing, for each off-target sites. Frequencies are presented as the average percentage of edited reads from HEK293T cells transfected with CBE-3 (n = 3).

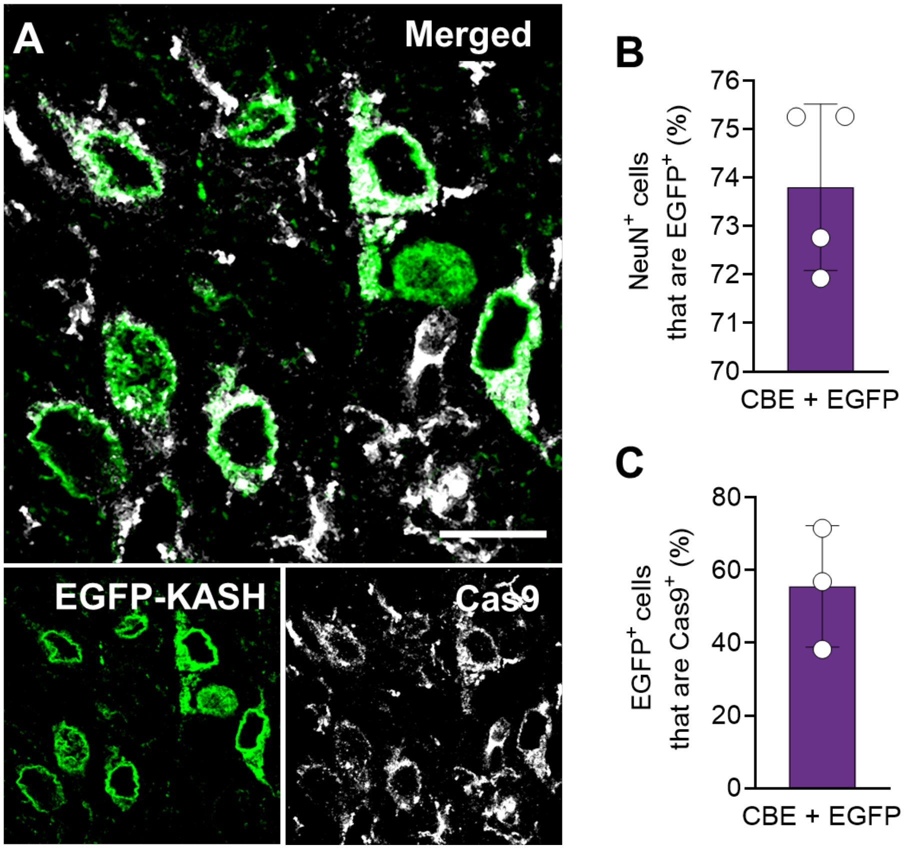


**Supplementary Figure S5. Striatal cells expressing EGFP-KASH also expressed the CBE protein. (A)** Representative immunofluorescence staining of the striatum four weeks after R6/2 mice were injected with 3 x $10^{10}$ particles each of dual AAV1 particles encoding the N- or C-terminal split-intein CBE domains and an additional 3 x $10^{10}$ particles of AAV1-EGFP-KASH. Scale bar; 15 µm. **(B)** Percentage of $NeuN^+$ cells that were $EGFP^+$ in the striatum of the injected R6/2 mice (n = 4). A total of 826 cells were counted. **(C)** Percentage of $EGFP^+$ cells that were $Cas9^+$ in the striatum of the injected R6/2 mice (n = 3). A total of 272 cells were counted. (**B** and **C**) Bars represent means and error bars indicate S.D.

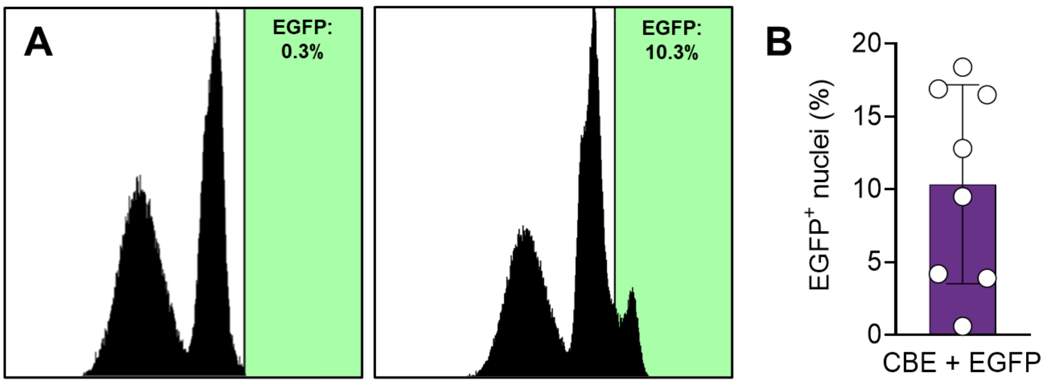


**Supplementary Figure S6. FACS enrichment of EGFP$^+$ nuclei from striatal tissue. (A)** Representative FACS histograms illustrating EGFP fluorescence intensities of nuclei isolated from striatal tissue of (left) control and (right) R6/2 mice four weeks after injection of 3 x $10^{10}$ particles each of dual AAV1 particles encoding the N- or C-terminal split-intein CBE domains and 3 x $10^{10}$ particles of AAV1-EGFP-KASH. **(B)** Quantitation of the percentage of EGFP$^+$ nuclei isolated from striatal tissue (n = 8). Unstained nuclei from an uninjected mouse was used as a negative control for gating. Bar represents mean and error bar indicates S.D.

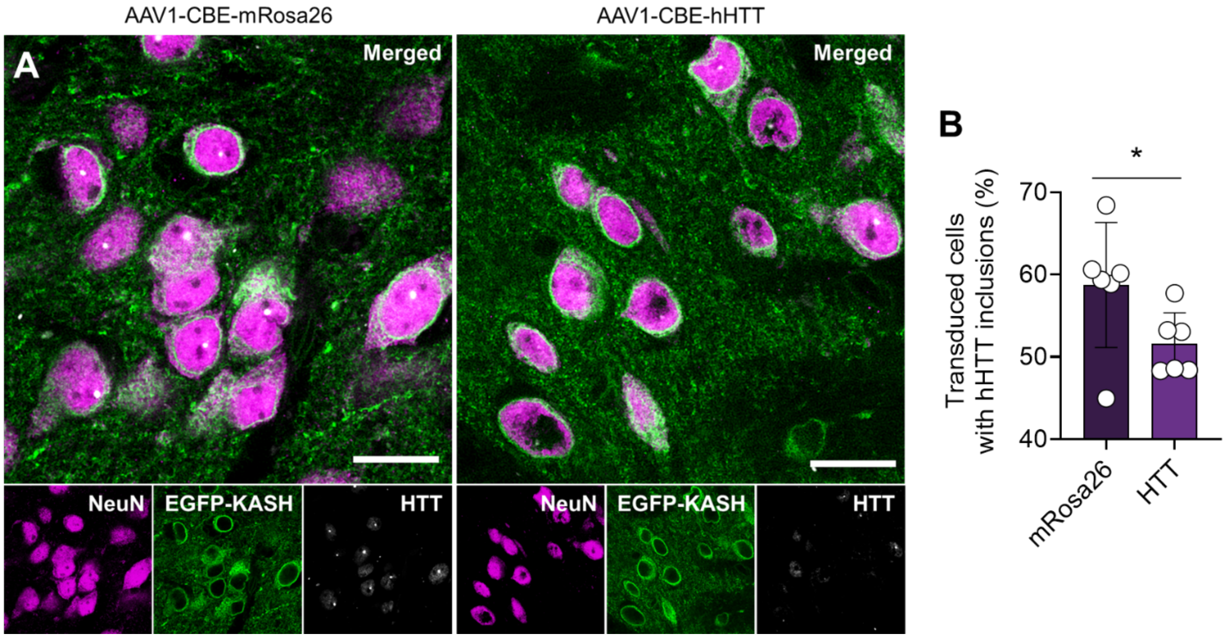


**Supplementary Figure S7. Base editing reduced the abundance of mutant HTT (mHTT) immunoreactive inclusions in the striatum of R6/2 mice**. **(A)** Representative immunofluorescence staining of the striatum four weeks after R6/2 mice were injected with 3 x $10^{10}$ particles each of dual AAV1 particles encoding the N- or C-terminal split-intein CBE domains and 3 x $10^{10}$ particles of AAV1-EGFP-KASH. Scale bar; 15 µm. **(B)** Quantitation of the percentage of transduced cells with visible HTT immunoreactive inclusions within the striatum of injected R6/2 mice. A total of >100 cells were counted per animal (n = 6). Bars represent means and error bars indicate S.E.M. *P < 0.05; one-tailed unpaired t-test.

### REFERENCES


1. Hsu, P.D., Scott, D.A., Weinstein, J.A., Ran, F.A., Konermann, S., Agarwala, V., Li, Y., Fine, E.J., Wu, X., Shalem, O. *et al.* (2013) DNA targeting specificity of RNA-guided Cas9 nucleases. *Nat. Biotechnol.*, **31**, 827-832.
